# The cell-wall-localised BETA-XYLOSIDASE 4 contributes to immunity of Arabidopsis against *Botrytis cinerea*

**DOI:** 10.1101/851170

**Authors:** Athanas Guzha, Robert McGee, Denise Hartken, Patricia Scholz, Daniel Lüdke, Kornelia Bauer, Marion Wenig, Krzysztof Zienkiewicz, Ivo Feussner, A. Corina Vlot, Marcel Wiermer, George Haughn, Till Ischebeck

## Abstract

Plant cell walls constitute physical barriers that restrict access of microbial pathogens to the contents of plant cells. The primary cell wall of multicellular plants predominantly consists of cellulose, hemicellulose and pectin. In Arabidopsis, a cell wall-localised protein, BETA-XYLOSIDASE 4 (BXL4) that belongs to a seven-member BETA-XYLOSIDASE (BXL) gene family was induced upon infection with the necrotrophic fungal pathogen *Botrytis cinerea* and mechanical wounding in a jasmonoyl isoleucine (JA-Ile) dependent manner. Ectopic expression of the BXL4 gene in Arabidopsis seed coat epidermal cells was able to rescue a *bxl1* mutant phenotype suggesting that like BXL1, BXL4, had both xylosidase and arabinosidase activity and acts *in mura* on cell wall polysaccharides. *bxl4* mutants show a compromised resistance to *B. cinerea.* Upon infection, *bxl4* mutants accumulated reduced levels of JA-Ile and camalexin. Conditional overexpression of *BXL4* resulted in enhanced expression of *PDF1.2* and *PAD3* transcripts both before and after *B. cinerea* infection. This was associated with reduced susceptibility of the transgenic lines to *B. cinerea*. These data suggest that remodelling or degradation of one or more cell wall polysaccharides is important for plant immunity against *B. cinerea* and plays a role in pathogen-induced JA-Ile and camalexin accumulation.

**One-sentence summary:** BXL4 is a putative bifunctional xylosidase/arabinofuranisodase localising to the apoplast, important for immunity against the necrotrphic pathogen *B. cinerea*.

## INTRODUCTION

Plants are continuously exposed to a plethora of biotic threats such as herbivorous insects and microbial pathogens. To help mitigate pathogen threats, plants have evolved various inducible and constitutive defence mechanisms against the different biotic and abiotic perturbations (Schutzendubel and Polle, 2002; Jones and Dangl, 2006). Induced immune responses are diverse and include the production of various phytohormones that are activated by different classes of microbial pathogens, insect pests and abiotic stresses (McDowell and Dangl, 2000). Important constitutive barriers that plant pathogens must overcome to access cellular contents are the cuticle and the plant cell wall (Vorwerk et al., 2004; Underwood, 2012; Engelsdorf et al., 2017). The importance of cell walls for plant immunity is demonstrated by the abundance of cell wall degrading enzymes (CWDEs) that microbial pathogens secrete in order to successfully invade plant tissues (Glass et al., 2013; Quoc & Bao Chau, 2017). Plant cell walls consist of a complex meshwork of polysaccharides, where cellulose microfibrils are cross-linked by various hemicelluloses and embedded in a pectic matrix. Cellulose consists of (1-4)-β-linked D-glucose residues and is synthesised by cellulose synthase complexes located in the plasma membrane (Somerville, 2006; Carpita, 2011). Hemicelluloses are a diverse group of polysaccharides. In Arabidopsis, the most abundant hemicellulose is xyloglucan, which is characterised by a (1,4)-β-linked glucan regularly substituted with (1-6)-α-xylosyl residues (Liepman et al., 2010; Scheller and Ulvskov, 2010; Höfte and Voxeur, 2017).

Pectin is the most complex cell wall polysaccharide and its biosynthesis involves at least 67 different enzyme activities (Harholt et al., 2010). Pectin consists of four types of polysaccharides: homogalacturonan (HG), xylogalacturonan and rhamnogalacturonan (RG) I and II (Mohnen, 2008). Homogalacturonan, xylogalacturonan, and RG-II are all characterised by the presence of a (1-4)-α-D-galacturonic acid backbone, whereas RG-I, has a backbone alternating in (1-2)-α-L-rhamnose and (1-4)-α-D-galacturonic acid residues (O’Neill et al., 1990; Ridley et al., 2001; Mohnen, 2008; Mohnen, et al., 2008). RG-I is also characterised by arabinan, galactan and arabinogalactan side chains, and xylan side chains were recently proposed to be present as well (Ralet et al., 2016). In Arabidopsis, pectin is the most abundant polysaccharide in the primary cell wall, and it is important for the regulation of cell wall mechanical properties during growth and development. It also influences water imbibition of seeds, pollen tube growth, leaf and flower abscission, fruit ripening, and cell wall integrity induced signalling (Mohnen, 2008; Arsovski et al., 2009; Harholt et al., 2010; Kohorn and Kohorn, 2012).

In addition to polysaccharides, plant cell walls also contain various proteins (Albenne et al., 2014). These proteins comprise only a small proportion of plant cell wall components, but they are an integral part of the cell wall as they contribute to its structural integrity, or modify cell wall composition during plant development and in response to environmental cues (Sommer-Knudsen et al., 1998; Fry, 2004; Passardi et al., 2004). Modifications of pectin such as acetylation and methylesterification (Liu et al., 2018) are known to play a role in cell wall integrity maintenance and resistance to pathogens. For example, Arabidopsis *rwa2* (*reduced wall acetylation 2*) mutants, that have reduced pectin acetylation, are more resistant to *B. cinerea* (Manabe et al., 2011). Pectin methylesterase inhibitors (PMEIs) inhibit the activity of endogenous Arabidopsis pectin methylesterases (PMEs), which remove methylesters present on HG (Willats et al., 2001). Arabidopsis plants overexpressing PMEI-1 and PMEI-2 show enhanced resistance to the phytopathogens *B. cinerea* and *Pectobacterium carotovorum* (Lionetti et al., 2007). In Arabidopsis, four members of a berberine bridge enzyme-like family were found to be responsible for oxidation of oligogalacturonides (OGs) derived from homogalacturonan hydrolysis. Although the oxidised OGs trigger weaker immune responses, they are more resistant to hydrolysis by *B. cinerea* enzymes. Accordingly, Arabidopsis plants overexpressing these oxidases are less susceptible to this pathogen (*Benedetti et al., 2018*). However, given the various modifications that occur in pectin, it remains to be fully elucidated how they impact plant pathogen interactions. It has also been suggested that the activity of microbe derived polygalacturonases in Arabidopsis results in the generation of oligogalacturonide fragments derived from the hydrolysis of HG, that act as damage associated molecular patterns (DAMPs) and trigger cell wall damage signalling, thus, enhancing disease resistance (Ferrari et al., 2008; Gravino et al., 2017; Benedetti et al., 2018).

As highlighted above, most defence mechanisms involving pectin are attributed to homogalacturonan together with its derivatives, and the contribution of RG-I to plant defence is relatively unknown. In this study, we investigated the role of the Arabidopsis protein BXL4 (BETA-D-XYLOSIDASE 4, AtBXL4, XYL4) in the modification of RG-I and its impact on plant immunity. All seven Arabidopsis BXL family members (termed as BXL1-BXL7) possess predicted glycosyl hydrolase domains whilst some have predicted signal peptides for extracellular localisation (Goujon et al., 2003). Only BXL1 has a known function and was shown to be a bifunctional β-D-xylosidase/α-L-arabinofuranosidase (Goujon et al., 2003; Minic et al., 2004; Arsovski et al., 2009). BXL1 activity is important for extrusion of mucilage upon hydration of Arabidopsis seeds (Arsovski et al., 2009) and is associated with tissues undergoing secondary cell wall thickening (Goujon et al., 2003). We provide evidence that BXL4 localises to the plant cell wall, where it acts on both xylans and arabinans. We show that *BXL4* expression is induced by *B. cinerea* infection and mechanical wounding in a jasmonoyl-isoleucine (JA-Ile) dependent manner. Accordingly, *bxl4* mutants are more susceptible when challenged with *B. cinerea* and display reduced levels of JA-Ile and camalexin after infection. Conversely, overexpression of *BXL4* results in increased transcript accumulation of the *B. cinerea*-responsive marker genes *PDF1.2* and *PAD3* and enhanced resistance to *B. cinerea*.

## RESULTS

### BXL4 expression is induced by wounding and B. cinerea infection in a jasmonoyl-isoleucine-dependent manner

According to publicly available databases, the expression of *BXL4* is upregulated by infection with various pathogens (Winter et al., 2007; Hruz et al., 2008). To confirm that *BXL4* is a stress-induced gene, its expression pattern after mechanical wounding and infection with *B. cinerea* was analysed (Figure 1). Generally, *BXL4* gene expression is relatively low in Col-0 grown under normal conditions (Supplemental figure 1). However, the *BXL4* gene expression was induced 16-fold upon mechanical wounding of the rosettes of Col-0 (Figure 1A). Induction of *BXL4* expression was also investigated in the jasmonoyl-isoleucine (JA-Ile) deficient mutant line *dde2-2* (von Malek et al., 2002), because JA-Ile regulates the expression of many wounding responsive genes (Howe et al., 2018). Relative to wild-type expression, upregulation of *BXL4* transcript levels after wounding was greatly reduced in the *dde2-2* mutant (Figure 1A). Because Arabidopsis defence against necrotrophic pathogens is associated with JA-Ile (Glazebrook, 2005; Pieterse et al., 2012), accumulation of *BXL4* transcript after infection with *B. cinerea* was quantified. There was a significant 12 to 20-fold induction of *BXL4* transcript accumulation in Arabidopsis 72 hours post inoculation (hpi) with *B. cinerea* (Figure1B, C). To investigate whether the induction of *BXL4* expression after *B. cinerea* infection also occurs in unchallenged systemic tissues, the *BXL4* expression was analysed in distal leaves after a local *B. cinerea* drop-inoculation. This analysis revealed that the *BXL4* expression is induced in the uninfected distal leaves as well (Figure 1C).

**Figure 1:**
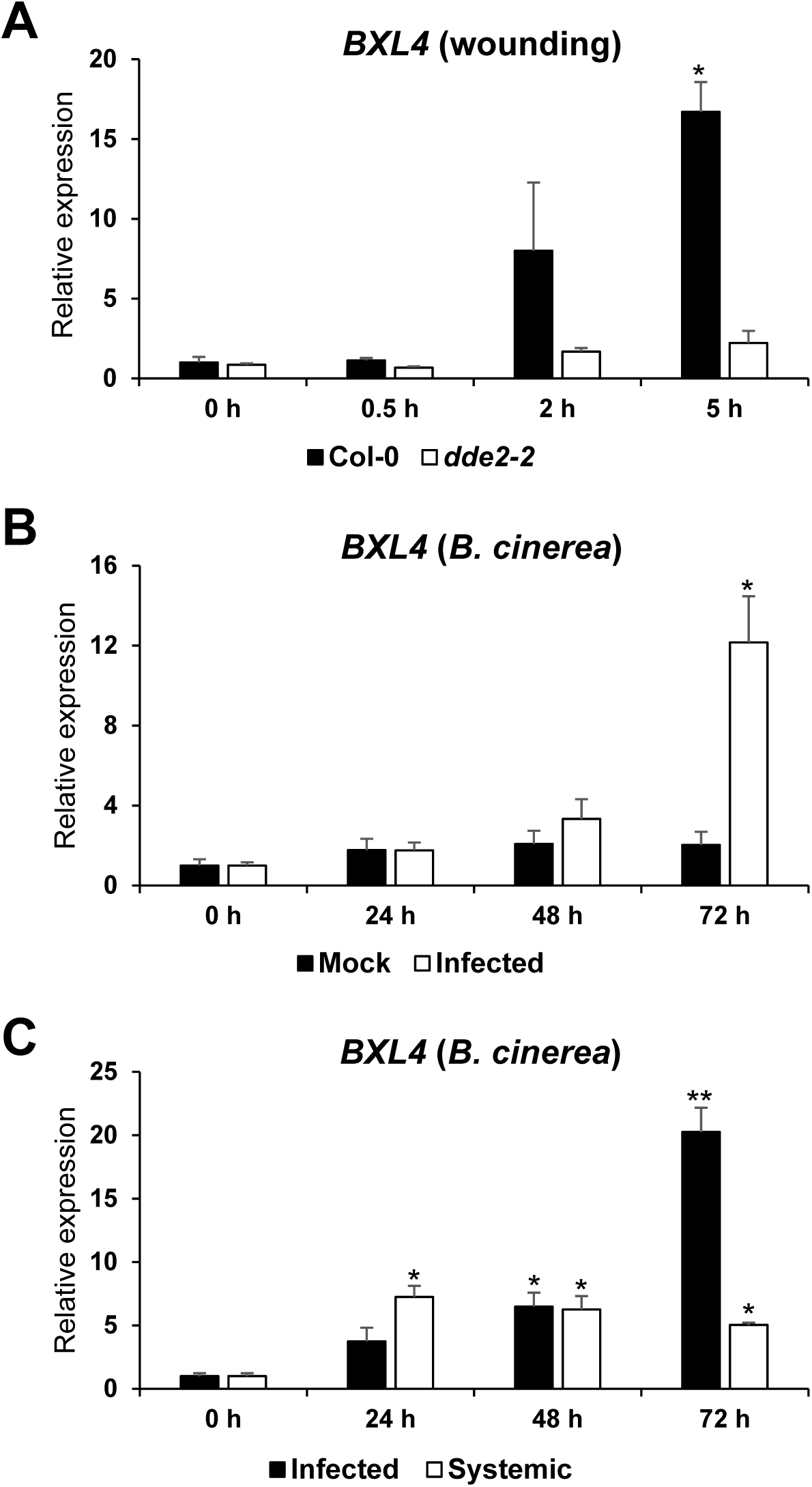
*BXL4* is induced upon mechanical wounding and *B. cinerea* infection in a JA-Ile-dependant manner. **(A)** Expression pattern of *BXL4* in Col-0 and the JA-Ile-deficient mutant *dde2-2.* RNA extracted from 6 weeks old plants before wounding (0 h) and at 0.5, 2 and 5 h after wounding. **(B)** Relative expression of *BXL4* in leaves of 6 week-old Col-0 plants after leaves were drop inoculated with *B. cinerea* conidiospores suspension or Vogels buffer (mock). **(C)** Relative expression of *BXL4* in infected or unchallenged systemic (distal) leaves at 0 (before inoculation), 24, 48 and 72 h after inoculation. (n=3 biological replicates), statistical difference to WT 0 h (Student‘s *t*-test; *p<0.05, **p<0.01). Experiments were conducted three times with similar results.

### BXL4 localises to the apoplast

Cell wall modification *in mura* is one important aspect of plant defence (Ferrari et al., 2012; Lionetti et al., 2017). The Arabidopsis protein BXL4, annotated as beta-xylosidase, possesses a predicted signal peptide for secretion to the apoplast (Goujon et al., 2003) and showed increased abundance in the Arabidopsis apoplast after *P. syringae* and *B. cinerea* infection (Breitenbach et al., 2014; Sham et al., 2014). To confirm cell wall localization, *BXL4*-*CITRINE* was stably expressed in seed coat epidermal cells, where the apoplast can be easily distinguished from the plasma membrane as it forms mucilage pockets (Haughn and Western, 2012; Tsai et al., 2017). A *BXL4* construct with a C-terminal CITRINE fusion (*BXL4*-*CITRINE*) driven by the strong, seed coat specific *TBA2* (*TESTA ABUNDANT2*) promoter (Tsai et al., 2017; McGee et al., 2019) was generated and transformed into Arabidopsis Col-0 wild-type plants. Arabidopsis seed coat epidermal cells of T2 transgenic seeds expressing *pTBA2:BXL4*-*CITRINE* in Col-0 were visualised under a confocal microscope 7 d post anthesis. The BXL4-CITRINE fluorescence could be detected in the mucilage pocket (Figure 2).

**Figure 2:**
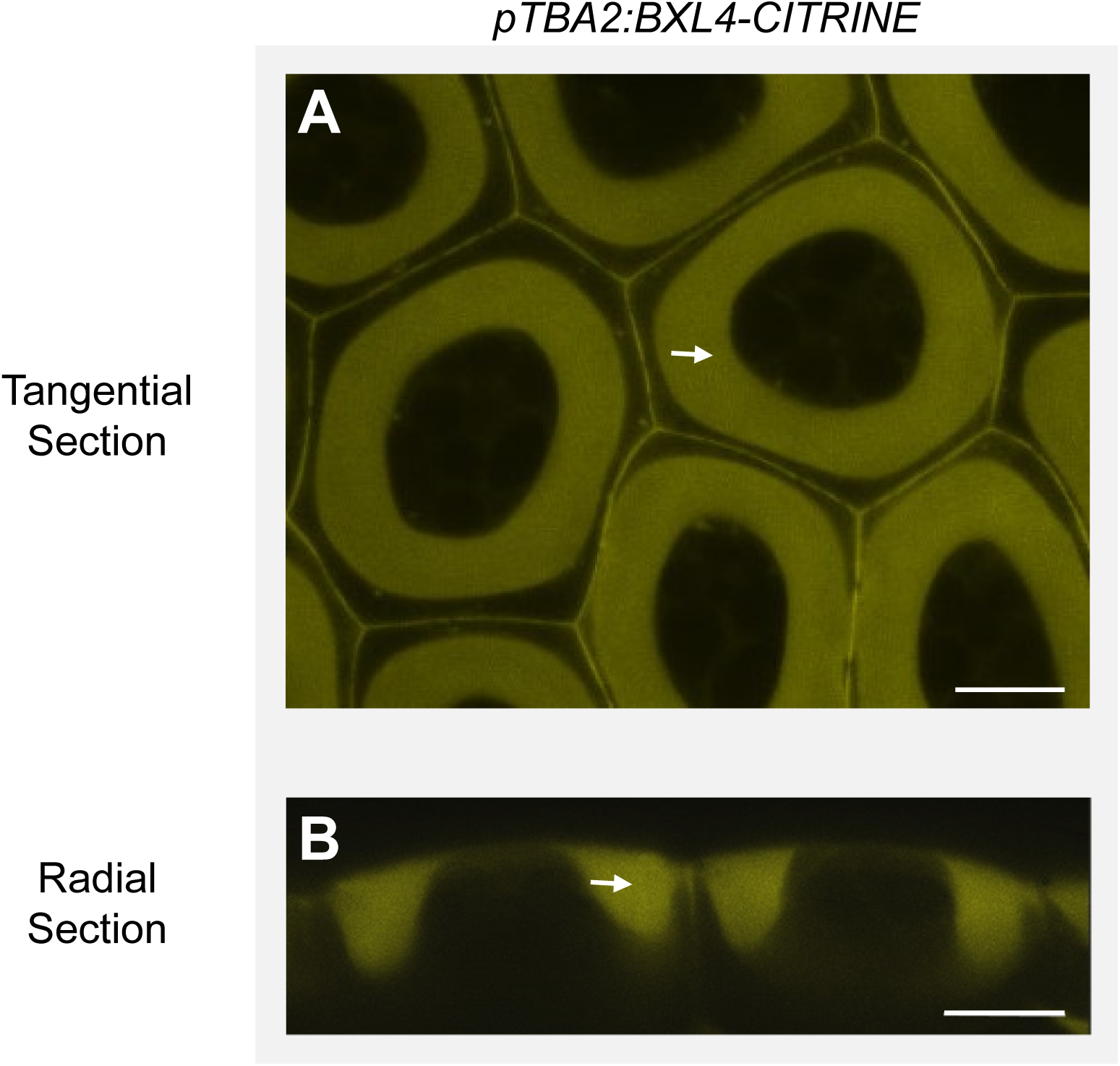
BXL4-CITRINE localises to the apoplast in Arabidopsis Col-0 seed coat epidermal cells. BXL4-CITRINE stably expressed under the control of the TBA2 promoter localises to the mucilage pocket (see arrows), a section of the apoplast of Arabidopsis seed coat epidermal cells, at 7 days post anthesis. **(A)** Tangential confocal section. **(B)** Radial confocal section. Scale bars, 10 µm.

### BXL4 rescues the mucilage phenotype of bxl1 when expressed under the TBA2 promotor that drives expression in seed coats

Before investigating the physiological function of BXL4, we aimed to gain information on its enzymatic activity that has previously not been characterized. However, its close homolog, BXL1 (Supplemental figure 2), has been shown to have arabinosidase/xylosidase activity (Goujon et al., 2003; Minic et al., 2004). BXL1 is found in the mucilage pocket of seed coat epidermal cells (Tsai et al., 2017), where it modifies mucilage polysaccharides (Arsovski et al., 2009). The *bxl1* knockout mutant produces mucilage that extrudes much more poorly from the seed coat after hydration of the seeds in comparison to wild-type mucilage due to changes in its polysaccharide composition (Arsovski et al., 2009; Figure 3). Since BXL4 has not been identified in the seed mucilage (Tsai et al., 2017) and the *bxl4* knockout mutant lines had no obvious mucilage defect (Supplemental figure 3), we attempted to use complementation of the *bxl1* mucilage phenotype by expressing *BXL4* under a promoter that drives strong expression in the seed-coat. This way, we aimed to determine whether BXL4 has an enzymatic activity similar to that of BXL1. The *bxl1* knockout mutant was stably transformed with the *pTBA2*:*BXL4* construct with and without a C-terminal CITRINE tag. In addition, as positive controls, a *pTBA2-BXL1* construct with and without a C-terminal CITRINE tag was transformed into *bxl1*. T2 transgenic *bxl1* seeds containing either the *pTBA2*:*BXL1-CITRINE* or the *pTBA2*:*BXL4-CITRINE* construct showed apoplastic localisation of CITRINE in seed coat epidermal cells (Supplemental figure 4). The ability of all constructs to complement the *bxl1* mucilage extrusion defect was determined by placing T2 transgenic seeds in water and staining the mucilage with ruthenium red, a dye that stains acidic polysaccharides such as pectin (McFarlane et al., 2013). Sixteen out of 17 independent transformants carrying *pTBA2*:*BXL1* (Figure 3C and supplemental figure 4) and all 21 independent transformants carrying the *pTBA2*:*BXL4* (Figure 3D and supplemental figure 5) constructs with and without the CITRINE tag, showed normal mucilage extrusion. In contrast, BXL6, a BXL4 homolog (Supplemental figure 2) that does not possess a signal peptide for extracellular targeting (Based on SignalP-5.0 analysis; Armenteros et al., 2019) under control of the *TBA2* promoter failed to rescue the mucilage defect of *bxl1* (Supplemental figure 6). The Arabidopsis *bxl1* knockout mutant produces mucilage with a higher content of arabinose than the wild type (Arsovski et al., 2009). To investigate if the *bxl1* transgenic lines carrying *pTBA2*:*BXL4-CITRINE* had mucilage with a monosaccharide composition more similar to wild-type Ws than *bxl1*, we analysed the water-extracted mucilage from T2 seeds after mild shaking. Mucilage of Arabidopsis seeds is composed mainly of the pectin RG-I (Dean et al., 2007; Arsovski et al., 2009; Arsovski et al, 2010; Haughn & Western, 2012). Consistent with this, our monosaccharide analysis of the mucilage showed that rhamnose and galacturonic acid were the most abundant sugars (Supplemental figure 7). The *bxl1* mutant exhibited a 4-fold increase in abundance of arabinans in comparison to the wild type Ws, whereas the *bxl1* transgenic lines complemented with *pTBA2*:*BXL1-CITRINE* or *pTBA2*:*BXL4-CITRINE* showed arabinose levels similar to the wild type (Figure 3E). Interestingly, the *bxl1* mutant line expressing *pTBA2*:*BXL4-CITRINE* showed a greater reduction in xylose levels compared to the wild-type control (Figure 3E).

**Figure 3:**
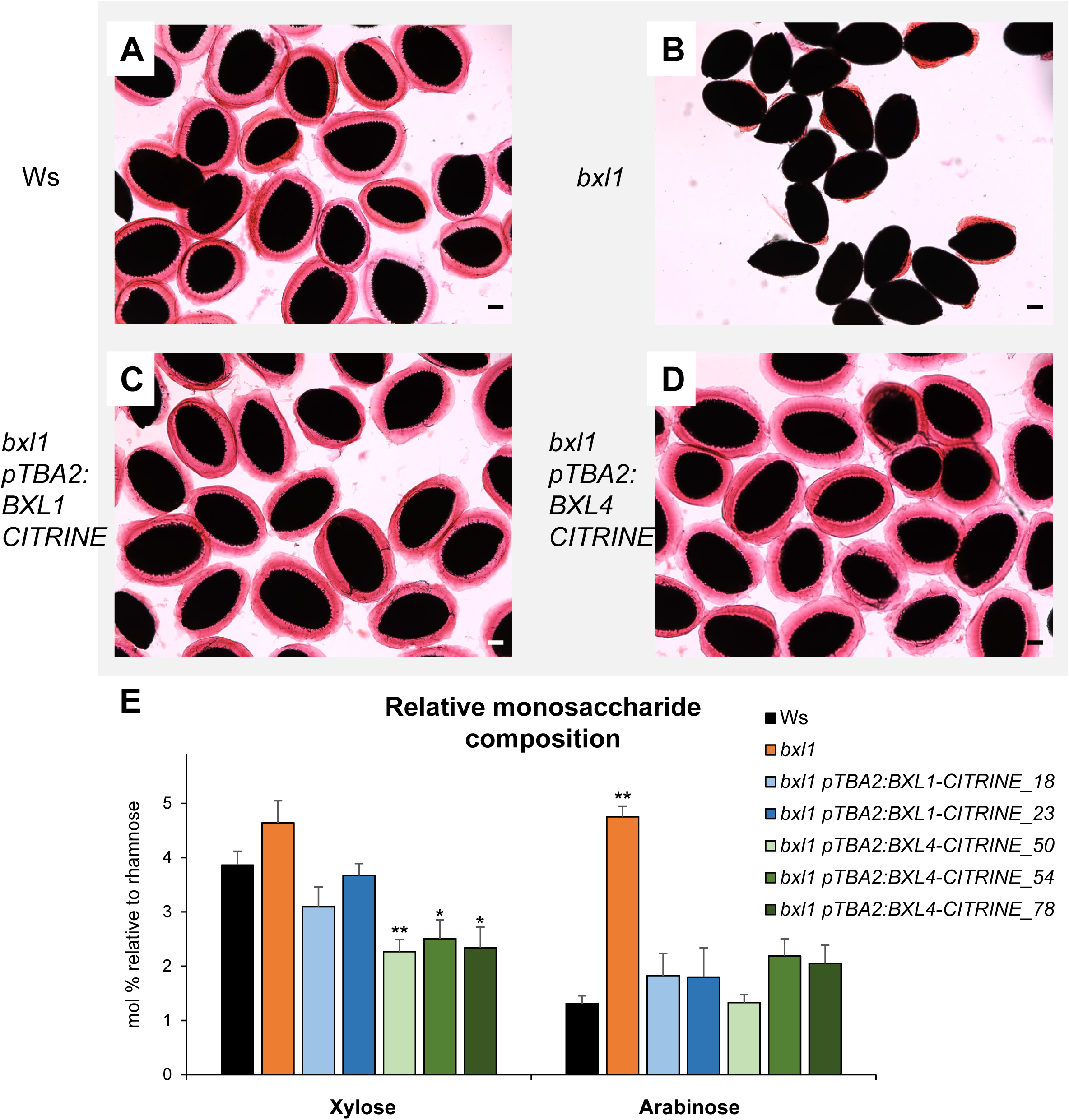
BXL4 can complement the mucilage phenotype of *bxl1*. In Ws wild-type plants, the extrusion of the mucilage forms a halo around the seeds that can be visualised by staining with ruthenium red **(A)**. *bxl1* seeds extrude their mucilage in a patchy manner **(B)**. The *bxl1* mucilage extrusion defect can be complemented by transgenic expression of *pTBA2*:*BXL1-CITRINE* **(C)** or *pTBA2*:*BXL4-CITRINE* **(D)**. Scale bars, 100 µm. **(E)** Relative monosaccharide composition of mucilage extracted from Ws, *bxl1, bxl1 pTBA2*:*BXL1-CITRINE* (line 18 and 23) and *bxl1 pTBA2*:*BXL4-CITRINE* (line 50, 54 and 78). Monosaccharide composition was determined by GC-MS and normalised to rhamnose. n=3 biological replicates. Error bars show SD, statistical difference to WT (Student‘s *t*-test); * indicates p<0.05, ** p<0.01

### Expression of BXL4 in Arabidopsis Col-0 seed coat epidermal cells results in less adherent mucilage

To enhance our understanding of the effects of BXL4 on Arabidopsis mucilage properties, and to test if the above observed putative xylosidase/arabinosidases activity could also be observed in wild-type Col-0 plants, Arabidopsis lines carrying the *pTBA2*:*BXL4* construct were made by transforming wild-type Col-0. The T2 seeds were shaken vigorously in water for 2 h before ruthenium red staining was done. The Col-0 lines with *pTBA2:BXL4* construct had a reduction in the volume of adherent mucilage compared to the wild-type control (Figure 4A-D). Indeed, quantification of the adherent mucilage volume using ImageJ v.1.84 (Schneider et al., 2012) revealed a significant reduction in the Col-0 lines transformed with the *pTBA2:BXL4* construct compared to the wild type Col-0 (Figure 4E). The mucilage from the T2 transgenic seeds was extracted and analysed for monosaccharide composition using GC-MS. Col-0 lines expressing *pTBA2:BXL4* had arabinose levels similar to the wild type, whilst the xylose composition was further depleted compared to Col-0 (Figure 4F). Together, these data suggest that BXL4 can complement the mucilage phenotype of the *bxl1* mutant and acts as a bifunctional β-D-xylosidase/α-L-arabinofuranosidase.

**Figure 4:**
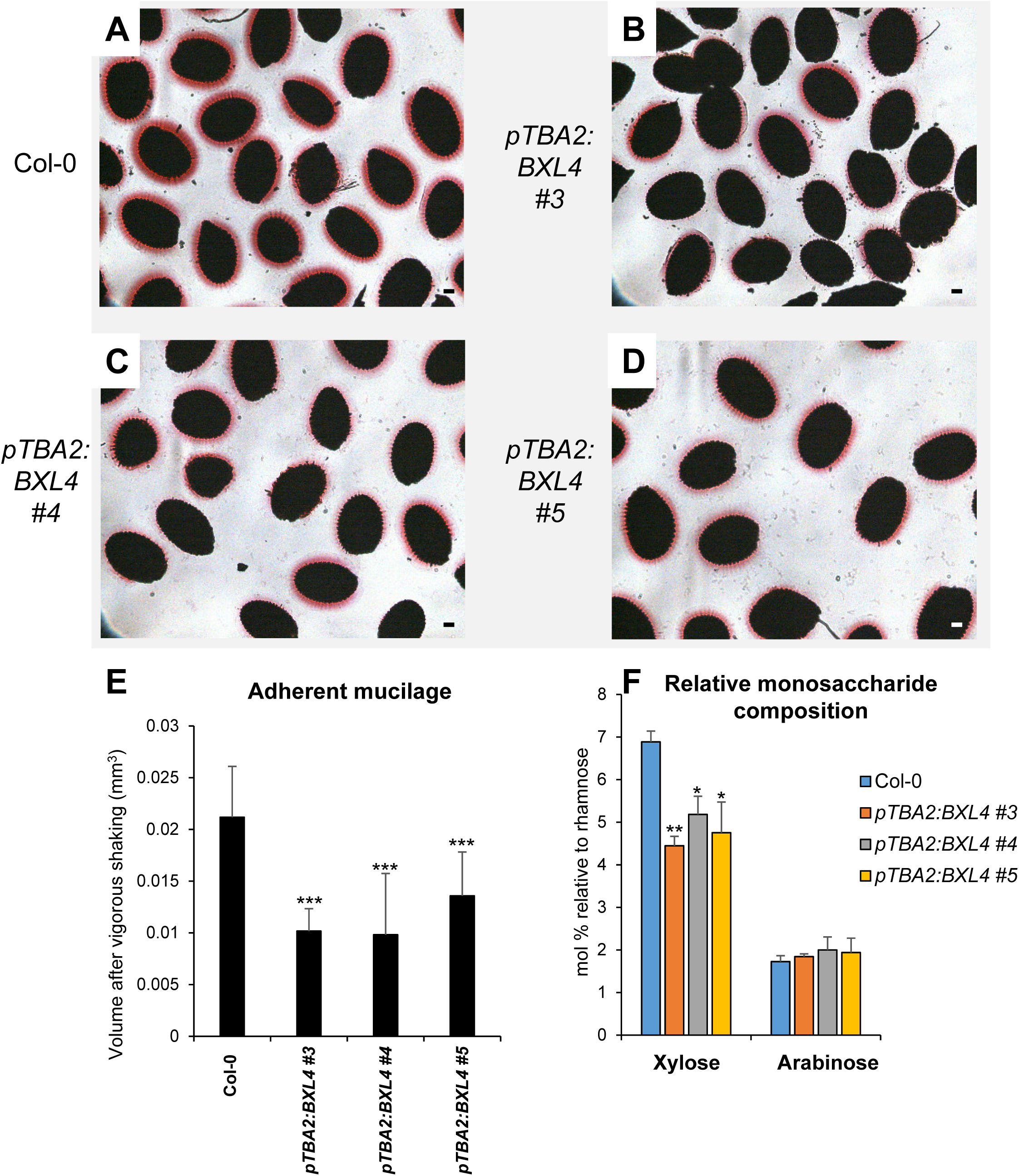
*BXL4* expression in Col-0 seed coat epidermal cells results in less adherent mucilage after vigorous shaking in water. Adherent mucilage of Col-0 **(A)**, *pTBA2*:*BXL4* in Col-0 line 3, 4 and 5 (**B**, **C** and **D** respectively) after vigorous shaking in water and staining with ruthenium red. Scale bar = 100 μm. **(E)** Quantification of the volume of adherent mucilage of Col-0, *pTBA2*:*BXL4* expression lines 3, 4 and 5 after vigorous shaking and staining with ruthenium red. n=10 seeds. **(F)** Monosaccharide composition of mucilage extracted from wild-type Col-0 and the three *pTBA2*:*BXL4* in Col-0 expression lines (3, 4 and 5). Monosaccharide composition was determined by GC-MS and normalised to rhamnose. n=3 biological replicates. Error bars show SD, statistical difference to WT (Student‘s *t*-test; * indicates p<0.05, ** p<0.01, *** p<0.001).

### Disruption of BXL4 does not affect plant growth

Alteration of the cell wall composition may impair normal growth and development of Arabidopsis plants (Noguchi et al., 1997). To test if the disruption of *BXL4* gene function affects the normal growth of Arabidopsis, two independent Arabidopsis T-DNA insertion lines, *bxl4-1* and *bxl4-2*, that carry insertions in exon 4 and 5, respectively, were obtained from the Nottingham Arabidopsis stock centre (Figure 5A). To confirm the position of the T-DNA insertion, the genomic DNA of the mutants was sequenced. In addition, *BXL4* transcript accumulation was analysed in the T-DNA insertion lines by quantitative reverse transcription polymerase chain reaction (qRT-PCR) using RNA extracted from 4-week old plants grown on soil and primers that flank the T-DNA insertion locus. Whereas we could not detect full-length *BXL4* transcripts using primers flanking the T-DNA insertion in the *bxl4-1* line, *BXL4* transcript accumulation was detectable, but significantly reduced, in *bxl4-2* plants (Figure 5B). Use of primers upstream of the T-DNA insertion site revealed a significant reduction of *BXL4* transcript accumulation in the *bxl4* mutants compared to Col-0 (Figure 5C and D). To evaluate if the disruption of *BXL4* gene function in *bxl4-1* and *bxl4-2* affects the monosaccharide composition of pectin in unchallenged tissues, water-extracted pectin from alcohol insoluble residues of leaf material from 6 weeks old Arabidopsis plants was analysed for monosaccharide composition using a modified method of gas chromatography-mass spectrometry (GC-MS) (Biswal et al., 2017). The monosaccharide compositions were normalised to rhamnose as its content is about proportional to the amount of RG-I (Saffer et al., 2017). Both *bxl4* mutants, in particular *bxl4-1*, showed an increased relative abundance of arabinose compared to Col-0 with the values not being significant for *bxl4-2*, though. In both lines, there was also no significant difference in the xylose content (Figure 5E) or any other pectin monosaccharide measured (Supplemental figure 8). Dot blot assays carried out on pectin extracted from the three genotypes also indicate in the *bxl4* mutants an increased abundance of long stretches of 1,5-linked arabinosyl residues that are known to be side chains of RG-I (Supplemental figure 9) (Arsovski et al., 2009). Despite these effects on leaf cell wall pectin composition, neither *bxl4* mutant exhibited any obvious growth defects and were comparable to Col-0 (Supplemental figure 10).

**Figure 5:**
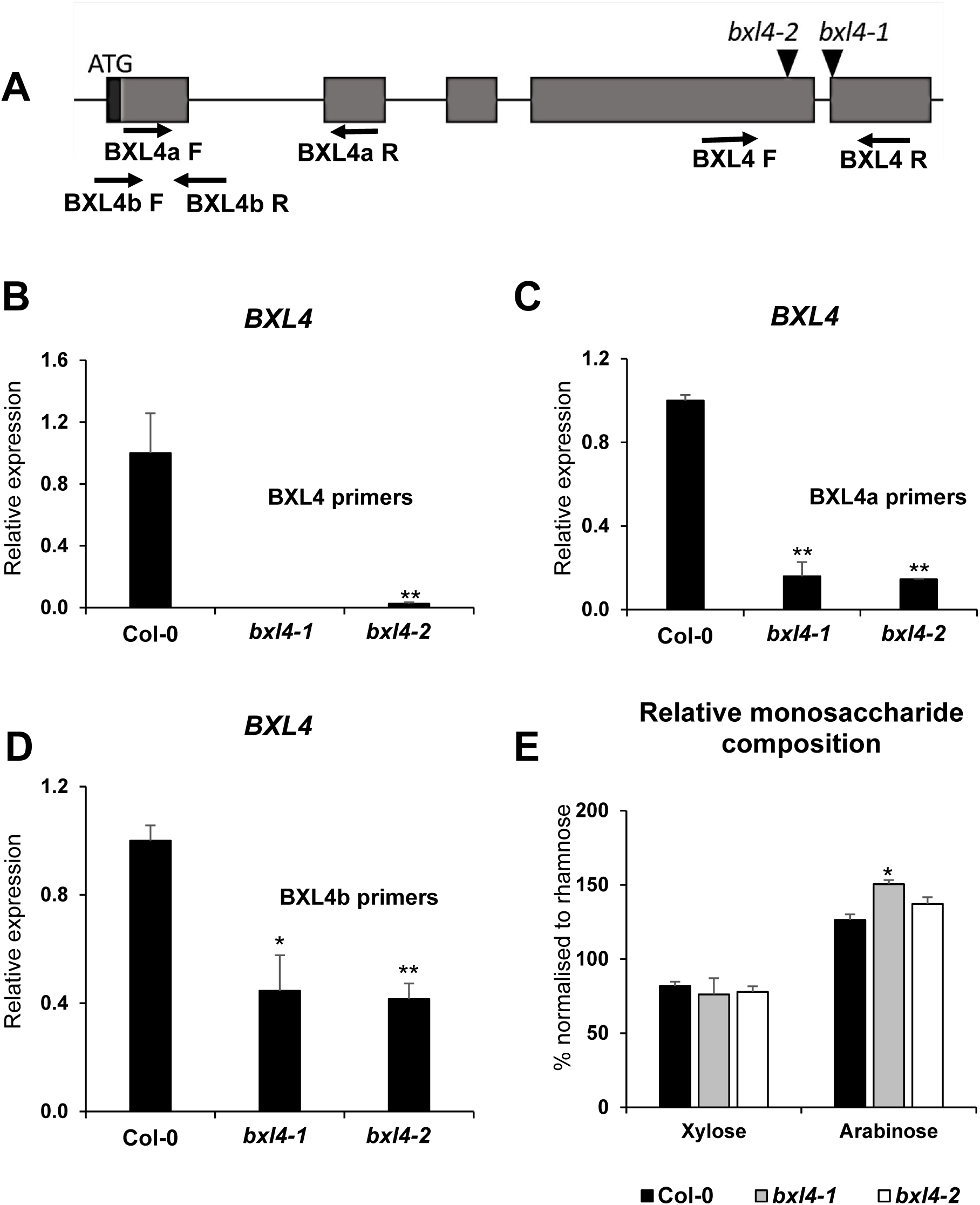
The disruption of *BXL4* has mild effects on the cell wall composition. **(A)** The intron-exon structure of *BXL4* and the positions of T-DNA insertions in *bxl4-1* and *bxl4-2.* The exons are represented by grey boxes, introns by black lines, black triangles show positions of the T-DNA insertions and the UTR is represented by a black box. Arrows indicate primers used for qRT-PCR analysis. **(B-D)** Relative expression of *BXL4* in leaves of WT and *bxl4* mutants as determined by qRT-PCR using primers shown in Supplementary table 1. Expression values were normalised to *ACTIN8* and are shown relative to Col-0. Error bars show SE (n=3 biological replicates), statistical difference to WT (Student‘s *t*-test) *p<0.05, **p<0.01. **(E)** Monosaccharide composition of water extracted pectin from Arabidopsis leaves of wildtype, *bxl4-1* and *bxl4-2* mutant lines. The monosaccharides were normalised to rhamnose. Extracted pectin was analysed by GC-MS. Error bars show SD (n=4 biological replicates), statistical difference to WT (Student‘s *t*-test; *p<0.05). Experiments were conducted three times with similar results.

### BXL4 contributes to resistance against B. cinerea

As the expression of *BXL4* is induced after *B. cinerea* infection (Figure 1), we tested if the *bxl4* mutants are compromised in resistance to *B. cinerea*. Therefore, plants were drop inoculated with *B. cinerea* conidiospore suspensions and the lesion area was measured 72 h post inoculation (Figure 6). The *bxl4* mutants developed significantly larger lesions compared to Col-0 (Figure 6A and C). The *mpk3* mutant was used as a control with enhanced susceptibility (Galletti et al., 2011). An additional method was used to quantify the disease susceptibility to *B. cinerea*, which involved spraying the plants with *B. cinerea* conidiospore suspension and quantifying the fungal β-ACTIN genomic DNA using qPCR (Ettenauer et al., 2014). Fungal DNA was quantified immediately after spraying (0 dpi) and 72 h after spray inoculation (3 dpi). The *bxl4* mutants showed a significantly higher abundance of fungal genomic DNA as compared to Col-0 (Figure 6B). To assess the effects of BXL4 overexpression on Arabidopsis defence reactions, three independent transgenic lines (OE1, OE2, and OE3) expressing *BXL4* in the Col-0 background under control of an estradiol-inducible promoter were generated and selected from seven independent transgenic lines. First, the expression of BXL4 was evaluated by qRT-PCR (Figure 6D and E) in these lines by investigating six week old Arabidopsis plants at 4 days after spraying with 0.01% Tween 20 (mock) or 50 µM β-estradiol. The mock treated plants did not show any induction of *BXL4* expression (Figure 6D), but there was significant induction of *BXL4* expression ranging from 15-60 fold increase compared to Col-0 in the β-estradiol induced lines (Figure 6E).

**Figure 6:**
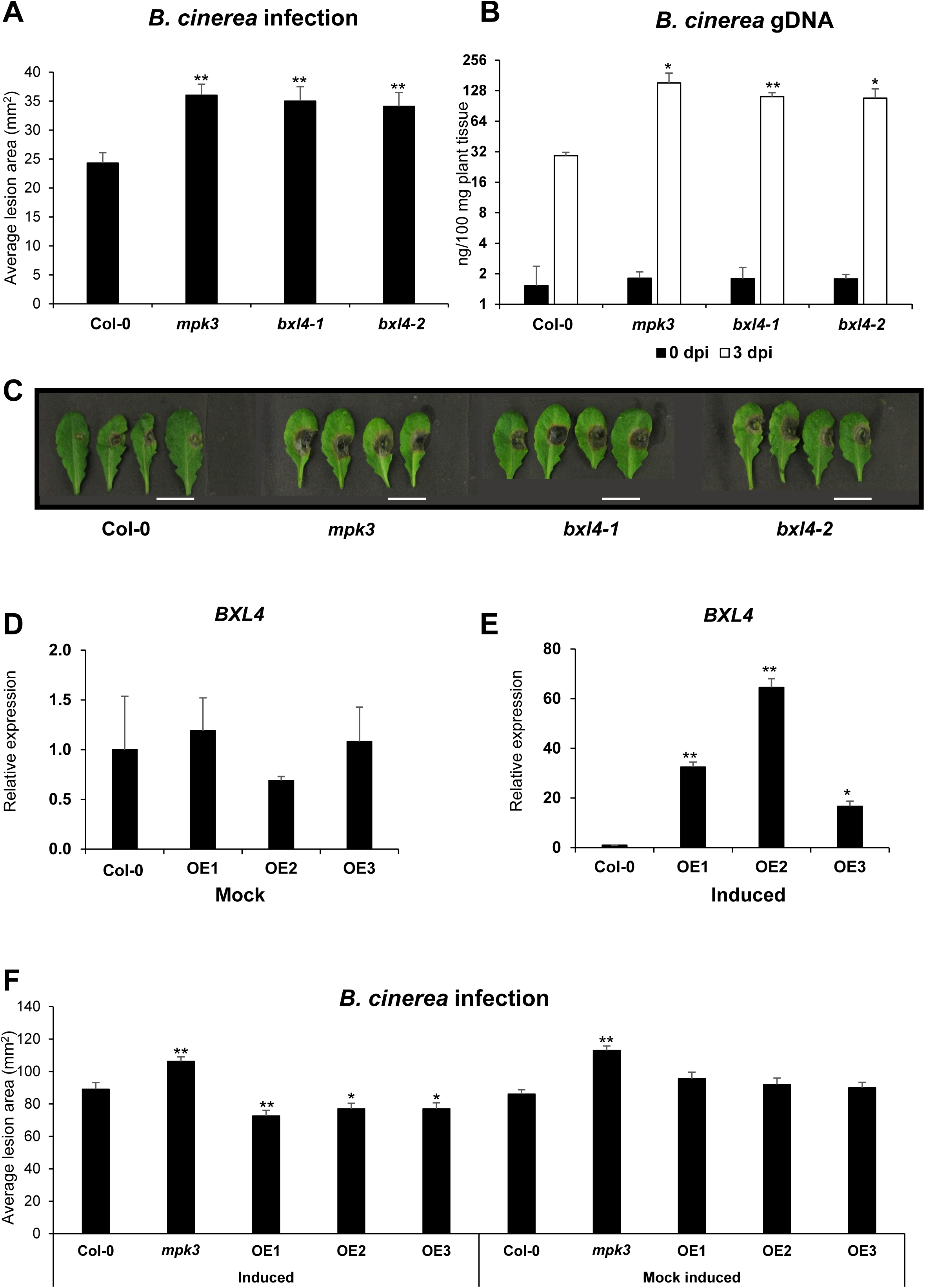
*bxl4* mutant lines are more susceptible to *B. cinerea* infection. **(A)** Infection phenotype of Col-0, *bxl4-1* and *bxl4-2* after *B. cinerea* infection. A minimum of 30 leaves from 5 independent plants were drop inoculated with 6 µl *B. cinerea* conidiospores, and lesion diameter measured with a digital caliper 3 days after infection and used to calculate lesion area. *mpk3* was used as the susceptible control. Error bars show SE (n ≥ 30 leaves), statistical difference to WT (Student‘s *t*-test), *p<0.05, **p<0.01. Experiment was conducted four times with similar results. **(B)** Infection phenotype measured after spraying plants with *B. cinerea* conidiospores and quantifying fungal genomic DNA with qPCR. Fungal genomic DNA was quantified immediately after infection (0 dpi) and 3 days after infection (3 dpi). Error bars represent SE (n=3 biological replicates), statistical difference to WT (Student‘s *t*-test), *p<0.05, **p<0.01. Experiment was conducted three times with similar results. **(C)** Lesion phenotype of detached leaves of Col-0, *mpk3*, *bxl4-1* and *bxl4-2* at 3 days post infection with *B. cinerea*. Scale bar = 10 mm. The relative expression of *BXL4* at 4 days after mock induction **(D)** and after β-estradiol induction **(E)** of *BXL4* inducible overexpression lines 1, 2 and 3 (OE1, OE2 and OE3). **(F)** Lesion area of Col-0, *mpk3* and inducible overexpression lines 1, 2 and 3. Plants were induced with β-estradiol for four days prior to infection with *B. cinerea* and lesion areas were scored 3 days after the infection. Error bars show SE (n ≥ 30 leaves), statistical difference to WT (Student‘s *t*-test), *p<0.05, **p<0.01.

We then tested if the overexpression of *BXL4* also had an effect on immunity to *B. cinerea*. The inducible *BXL4* overexpression lines were drop inoculated with *B. cinerea* conidiospores 4 days after β-estradiol induction, and lesion area was determined at 3 dpi. The *BXL4* overexpression lines developed smaller lesions compared to Col-0 (Figure 6F). In the mock-induced *BXL4* overexpression lines, there was no significant difference in lesion size compared to Col-0 (Figure 6F).

### Disruption of BXL4 alters the induction of plant defence associated genes upon mechanical wounding and B. cinerea infection

Mechanical wounding and attack of necrotrophic pathogens in Arabidopsis triggers defence responses, some of which are regulated via JA-Ile signalling (Howe et al., 2018). To test if the *bxl4* mutants are impaired in JA-Ile signalling, the transcript abundance of the JA-Ile marker genes *JAZ10* (Yan et al., 2007; Chung et al., 2008) and *PDF1.2 (Penninckx, 1998; Zarei et al., 2011*) was tested in mechanically wounded Col-0 and *bxl4* mutants (Figure 7). There was a significant induction of *PDF1.2* and *JAZ10* in Col-0 at 2 h post wounding, whereas the *bxl4* mutants had a significantly decreased transcript accumulation of *PDF1.2* (Figure 7A) and *JAZ10* (Figure 7B) at 2 h post wounding compared to Col-0. The transcript accumulation of the JA-Ile defence hormone associated marker genes *JAZ10* and *PDF1.2* were also evaluated after *B. cinerea* spray inoculation. The expression of *PDF1.2* was significantly induced in Col-0 at 24, 48 and 72 h post infection, whereas the *bxl4* mutants had expression that was compromised (Figure 7C). This trend was observed in three other independent experiments (Supplemental figure 11). Transcript accumulation of *JAZ10* showed a significant difference between the wild type and the *bxl4* mutants especially at 48 h post infection (Figure 7D). The relative expression of the *B. cinerea*-responsive marker gene *PAD3*, encoding an enzyme involved in antimicrobial camalexin biosynthesis, was also reduced significantly in *bxl4* plants especially at 48 h post inoculation compared to Col-0 (Figure 7E).

**Figure 7:**
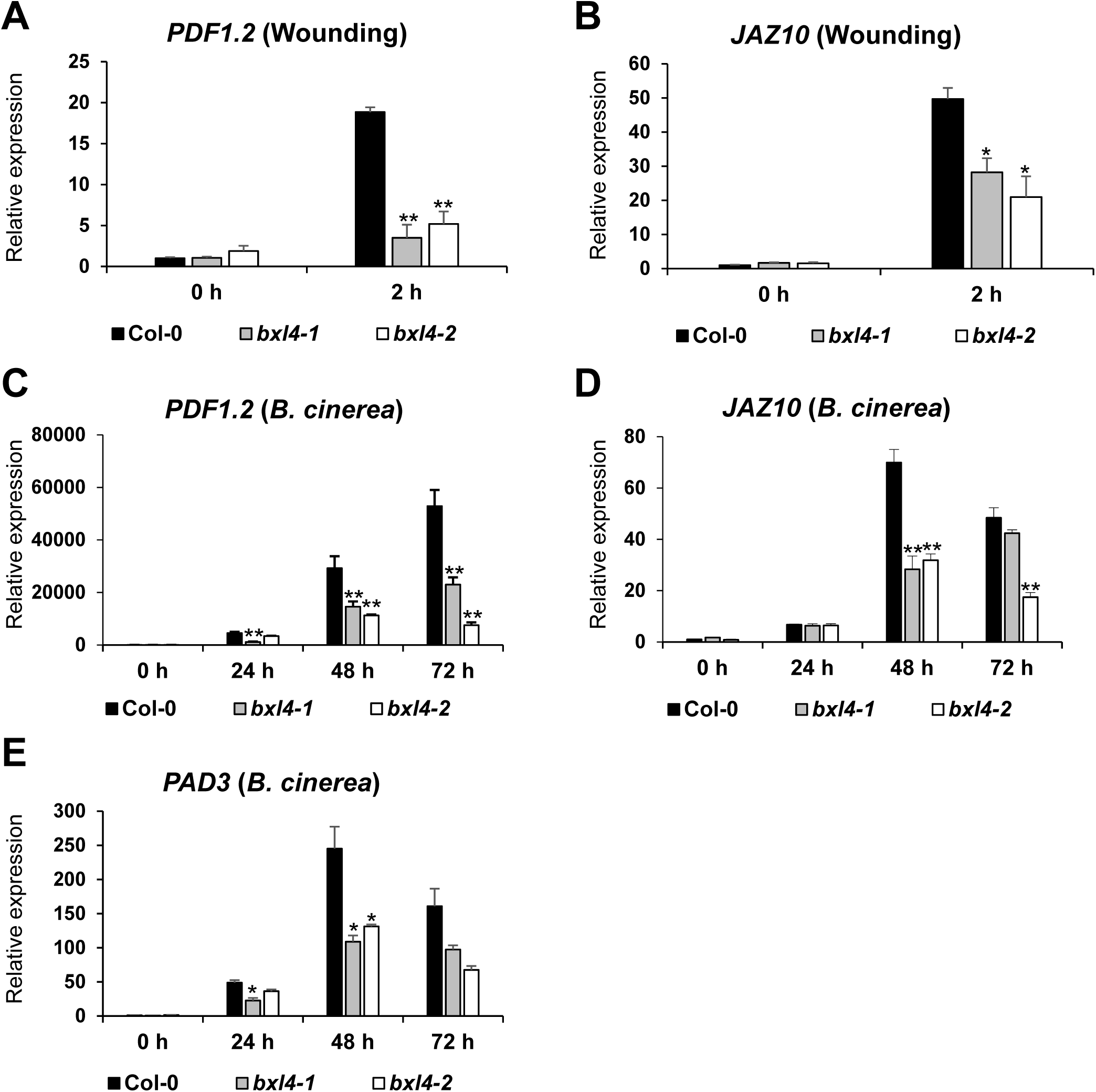
BXL4 acts upstream of jasmonoyl-isoleucine signalling upon wounding and *B. cinerea* infection. Relative expression of JA-Ile marker genes *PDF1.2* **(A)** and *JAZ10* **(B)** in Col-0, *bxl4-1*, and *bxl4-2* at 0 and 2 h post wounding. Relative expression of *PDF1.2* **(C)** and *JAZ10* **(D)** in 6 week old Col-0, *bxl4-1* and *bxl4-2* Arabidopsis plants at 0, 24, 48 and 72 h after infection with *B. cinerea.* **(E)** Relative expression of *PAD3* in *B. cinerea* infected Col-0 and *bxl4* mutant lines at 0, 24, 48, 72 h after infection. Expression values were normalised to the reference gene *ACTIN8* and are shown relative to the wild-type 0 h levels. Error bars show SE (n=3 biological replicates), statistical difference to WT (Student‘s *t*-test), *p<0.05, **p<0.01. Experiments were conducted three times with similar results

### bxl4 mutant plants have a reduced accumulation of JA-Ile and camalexin after infection with B. cinerea

We next assayed if the reduced expression of *PDF1.2* and *PAD3* in *bxl4* mutants correlates with a reduced accumulation of JA-Ile and camalexin upon infection with *B. cinerea* (Figure 8) (Ferrari et al., 2003, 2007; Scalschi et al., 2015; Nie et al., 2017). The *bxl4* mutants showed a slight reduction in JA-Ile abundance especially at 72 h post inoculation compared to Col-0 (Figure 8D). The abundance of camalexin after *B. cinerea* infection was reduced especially at 24 h post inoculation in the *bxl4* mutants compared to Col-0 (Figure 8F). There was no significant increase in the abundance of JA-Ile and camalexin after mock inoculation of the different genotypes. The wounded *bxl4* mutants also had a slightly reduced accumulation of JA-Ile compared to Col-0 (Supplemental figure 12). Other plant hormones or defence-related compounds did not show particular trends in *bxl4* plants after inoculation (Supplemental dataset 1).

**Figure 8:**
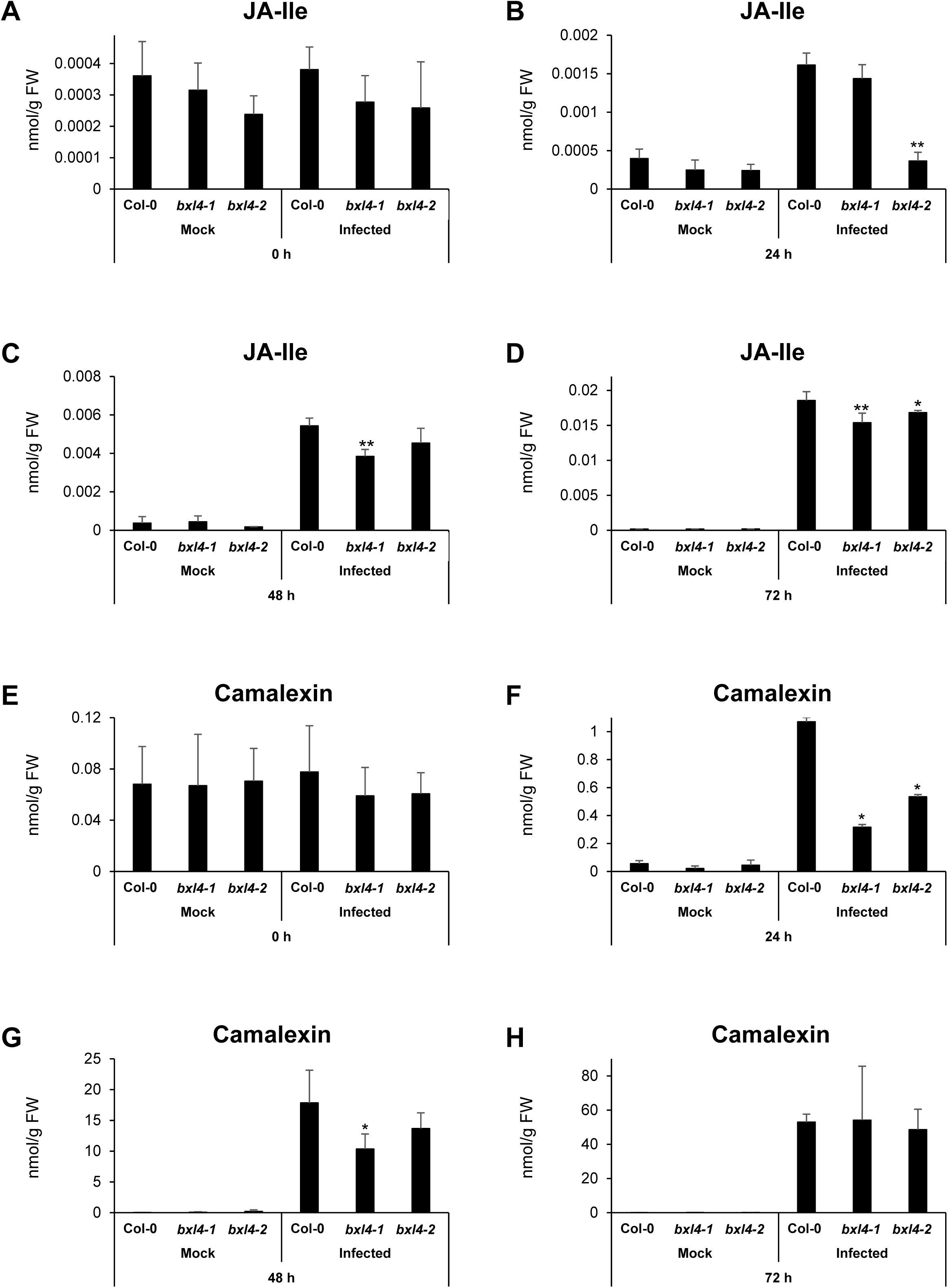
*bxl4* mutants have decreased accumulation of JA-Ile and camalexin upon infection with *B. cinerea*. Col-0 and *bxl4* mutant lines were spray induced with *B. cinerea* and the leaves were sampled at 0, 24, 48 and 72 h after infection. Extracted levels of JA-Ile **(A-D)** and camalexin **(E-H)** were analysed using nanoelectrospray coupled to a tandem mass spectrometer. Error bars represent standard deviation of 6 biological replicates, Statistical difference to WT (Student‘s *t*-test), *p<0.05, **p<0.01. Experiments were performed 3 times with similar results.

### BXL4 overexpression results in higher expression of PDF1.2 and PAD3 after B. cinerea infection

To test the effect of *BXL4* overexpression on the induction of *PDF1.2* and *PAD3* expression, the inducible *BXL4* overexpression lines were sprayed with β-estradiol or mock-induced by spraying 0.01% Tween20, and four days later the plants were infected with *B. cinerea.* Mock induction of the inducible lines did not result in any induction of *PDF1.2* or *PAD3* prior to infection (Figure 9A, C, time-point 0 h). Induction of *BXL4* by β-estradiol treatment, however, resulted in a strong upregulation of *PDF1.2* expression in the inducible overexpression lines compared to Col-0 (Figure 9B time point 0 h) prior to infection. In addition, a significant increase of *PAD3* expression was observed in one line (Figure 9D, time-point 0 h). The transcript accumulation of *PDF1.2* and *PAD3* was then evaluated at 24, 48 and 72 h post infection. The mock-treated overexpression lines plants did not show strong differences to the wild type (Figure 9A, C). When treated with β-estradiol four days prior to infection, however, differences could be observed. Here, the overexpression lines had a significantly increased *PDF1.2* transcript abundance at 24 h post infection (Figure 9B) and *PAD3* transcript was significantly higher especially at 48 and 72 h post infection (Figure 9D).

**Figure 9:**
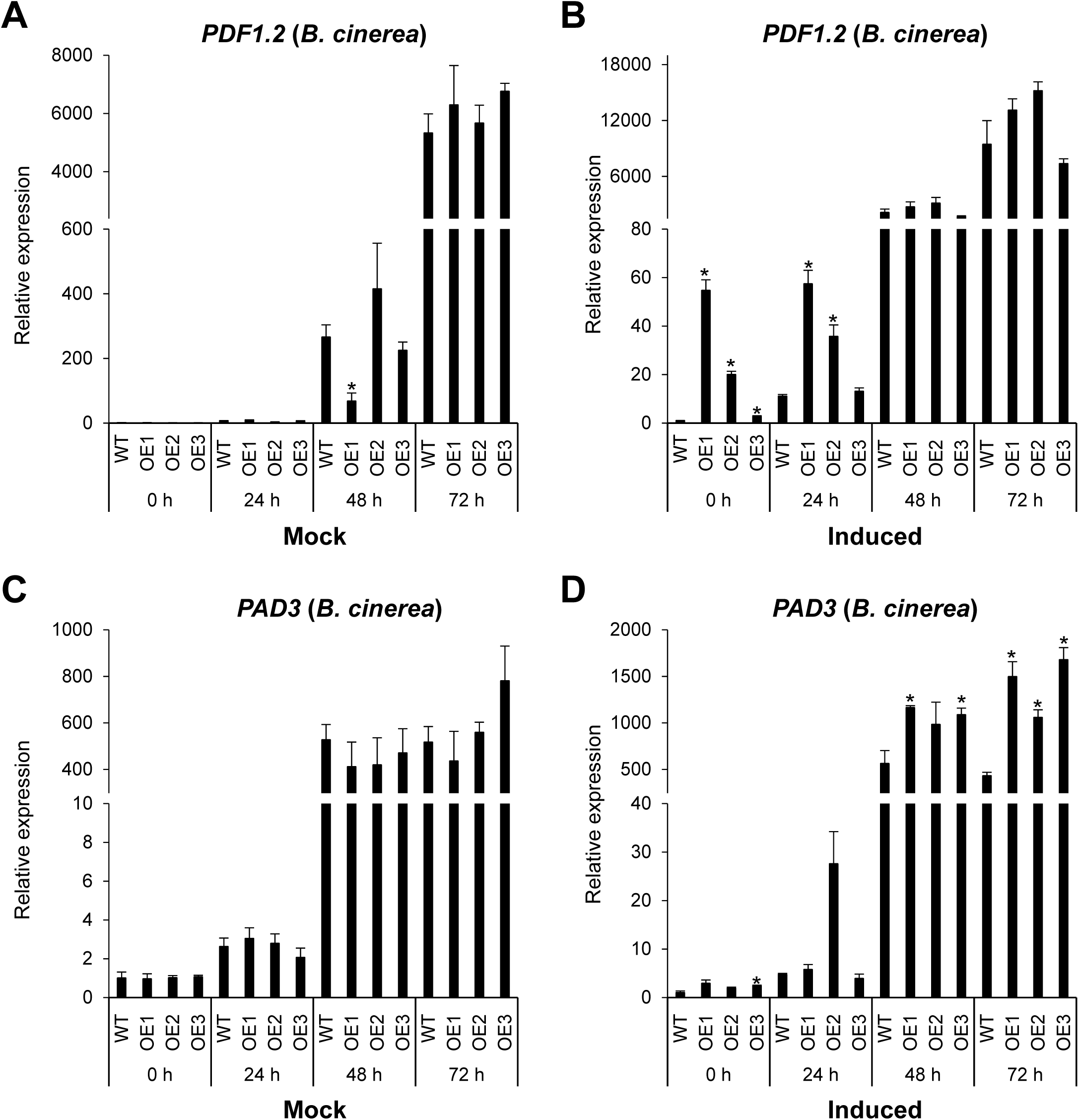
*BXL4* inducible overexpression lines have high induction of *PDF1.2 and PAD3*. Relative expression of *PDF1.2* after mock induction of *BXL4* **(A)** and β-estradiol induction of *BXL4* **(B)** in wild-type Col-0 and β-estradiol inducible *BXL4* overexpression lines 1, 2, and 3 (OE1, OE2 and OE3). *PAD3* expression measured in mock **(C)** and *BXL4* induced lines **(D)**. Expression was measured in leaf samples collected at 0 (4 days after *BXL4* induction and before *B. cinerea* inoculation), 24, 48, and 72 h after *B. cinerea* inoculation. Relative expression was measured by qRT-PCR, normalised to reference gene *ACTIN8*, and relative to the wild type 0 h. Error bars show SE of 3 biological replicates, statistical difference to WT (Student‘s *t*-test), *p<0.05, **p<0.01

## DISCUSSION

The importance of pectin modification in plant defences has been elucidated in several previous publications (Bethke et al., 2014; Lionetti et al., 2014, 2017) that highlighted the importance of HG methylation. Here, we provide evidence that BXL4 modifies another pectin polysaccharide, RG-I, and show that BXL4 contributes to immunity against *B. cinerea*.

### BXL4 likely acts as an arabinosidase and xylosidase

While our work was mostly concerned with the role of BXL4 in pathogen defence, we first investigated its molecular function in cell wall remodelling. The complexity of pectin is generated by a plethora of proteins with most of them being localized in the Golgi (Harholt et al., 2010). However, pectin can also be remodelled *in mura (Micheli, 2001; Bosch, 2005)* giving the plant more flexibility in the regulation of its cell wall architecture (Rui et al., 2018; Wu et al., 2018). Previous data on the cell wall proteome of Arabidopsis leaves expressing a *P. syringae* effector *AvrRpm1* indicated that BXL4 is such an enzyme acting *in mura* (Breitenbach et al., 2014). The apoplast localisation was also predicted by the algorithm of SignalP-5.0 (Armenteros et al., 2019), which indicated that the first 38 amino acids of BXL4 act a signal peptide for the secretory pathway. Our microscopic data confirms the apoplastic localization (Figure 2 and supplemental figure 4) making use of the seed coat mucilage system, where the cell wall can be easily distinguished from the plasma membrane (Haughn and Western, 2012; Tsai et al., 2017).

We also used the mucilage system for the *in vivo* investigation of the enzymatic activity of BXL4. BXL4 has a predicted glycosyl hydrolase domain (Goujon et al., 2003), and thus might be capable of carrying out similar enzymatic functions as its homolog BXL1 (57 % identity on the amino acid level). BXL1 modifies seed coat mucilage polysaccharides by reducing the amount of (1-5)-linked arabinans (Arsovski et al., 2009). Mutants with a loss of function mutation in *BXL1* have a distinct phenotype where seed mucilage has increased amount of arabinose and extrudes in a delayed and uneven manner when seeds are hydrated (Arsovski et al., 2009) (Figure 3B). BXL4, expressed in the seed coat epidermal cells, was able to complement this mutant phenotype (Figure 3D; Figure 3E) strongly suggesting that like BXL1, BXL4 has α-L-arabinofuranosidase activity. Expression of BXL4 under the strong *TBA2* promoter also decreased the amount of xylose in the pectinacious mucilage especially in the Col-0 background (Figure 4F). This also led to a reduced volume of adherent mucilage after vigorous shaking (Figure 4A-E), implying that the mucilage could be more loosely attached to the seed coat than in the wild type. This is consistent with the findings for *MUM5*, which encodes for a xylosyl transferase, and the mutant *mum5* with reduced xylose has mucilage loosely attached to the seed coat (Ralet et al., 2016). Recent findings by Ralet et al. (2016) suggest the presence of xylan side chains on RG-I that mediate the interaction with cellulose through non-covalent linkages. It is therefore conceivable that BXL4 acts on these xylan side chains and is therefore a bifunctional β-D-xylosidase/α-L-arabinofuranosidase. This bifunctional activity was also shown for BXL1 *in vitro* using proteins extracted from Arabidopsis wild-type and mutant stems (Goujon et al., 2003).

The increased levels of arabinans observed in the leaf cell walls (Figure 5E and supplemental figure 9) give further evidence that the arabinosidase activity of BXL4 acts on pectin, given that the cell wall polysaccharides were extracted using hot water and that hemicelluloses and cellulose are insoluble in water (Broxterman and Schols, 2018). However, the changes were relatively small considering the much bigger changes in cell wall composition in the mucilage of the *bxl1* mutant. The reason for this could be that *BXL4* is weakly expressed under regular growth conditions.

### BXL4 is required for full resistance to B. cinerea

The expression of *BXL4* in leaves is upregulated after wounding and after infection with *B. cinerea* (Figure 1). This upregulation is partially JA-Ile dependent (Figure 1C) similar to many genes involved in wound responses and pathogen defence (Howe et al., 2018). The induction of *BXL4* expression indicates that it plays a role in cell wall remodelling after wounding and pathogen attack. Cell wall remodelling has been previously described to occur after both these stresses. For example, the degree of pectin methylesterification is altered in Arabidopsis in response to attack from fungal pathogens (Lionetti et al., 2012). In many interactions between plants and pathogens it was noted that a high degree of methylesterification results in reduced susceptibility of the plants to pathogen as the pectin is more resistant to pectic enzymes (Lionetti et al., 2012, 2017; Liu et al., 2018). Wounding is also thought to trigger the induction of endogenous polygalacturonases that generate OGs important for defence responses (León et al., 2001). Similarly, we show genetic evidence for the involvement of *BXL4* in plant biotic stress tolerance, as the resistance to *B. cinerea* infection was compromised in the *bxl4* mutants (Figure 6A-C), while the conditional overexpression of *BXL4* resulted in decreased susceptibility to this pathogen (Figure 6F).

One way of action could be that cell wall modifications caused by BXL4 contribute to JA-Ile signalling processes, as the expression levels of the jasmonate regulated genes *PDF1.2* and *JAZ10* are influenced by both overexpression and downregulation of BXL4.

Another gene influenced by BXL4 and partially regulated by JA-Ile signalling is *PAD3* (Rowe et al., 2010) that catalyses the final step in camalexin production (Schuhegger et al., 2006). Camalexin has been shown to act as a phytoalexin not only against *B. cinerea* (Ferrari et al., 2007; Shlezinger et al., 2011) but also various other phytopathogens (Sanchez-Vallet et al., 2010; Schlaeppi et al., 2010). The reduction in the accumulation of camalexin in *bxl4* mutants, especially at early time-points after infection (Figure 8F), could be one of the factors that lead to enhanced susceptibility to *B. cinerea*. This estradiol inducible *BXL4* overexpression lines (OE1, OE2 and OE3), have a higher abundance of *PAD3* transcript especially at the later time points after *B. cinerea* infection (Figure 9D) and consequently they are more resistant to the pathogen.

Finally, JA-Ile also increases the expression of genes involved in JA-Ile synthesis generating a feed-forward loop (Farmer, 2007). Our data indicate that BXL4 could play a minor role in this feed forward loop, as JA-Ile levels were less strongly increased in the *bxl4* mutants at some time points after infection than in the wild type (Figure 8B-D).

The altered resistance in BXL4 knockdown and overexpression lines coming with changes in the expression levels of several marker genes raises the question how BXL4 could influence JA-Ile and defence signalling. Arabinan side chains that are likely trimmed by BXL4 play a role in cell wall architecture (Verhertbruggen et al., 2013). In Arabidopsis, the interspacing of homogalacturonan with arabinan-rich RG-I reduces crosslinking with Ca^2+^, thus making cell walls more flexible (Jones et al., 2003; Moore et al., 2008; Merced and Renzaglia, 2019). Trimming of arabinan side chains in wild-type plants could result in greater pectin crosslinking by Ca^2+^ as compared to the arabinan rich *bxl4* mutants, thus making the cell walls more recalcitrant to penetration by fungal hyphae.

In addition, the degradation of this cross-linked homogalacturan by polygalacturonases (Bellincampi et al., 2014; Sénéchal et al., 2014), would result in the formation of Ca^2+^ cross-linked oligogalacturonides, which elicit strong biological responses such as production of ROS, phytoalexins, callose and JA-Ile production (Kohorn and Kohorn, 2012; Bethke et al., 2014; Savatin et al., 2014; Mielke and Gasperini, 2019).

In conclusion, our study indicates that the trimming of RG-I side-chains by BXL4 is a factor of plant resistance.

## METHODS

### Plant and B. cinerea growth conditions

Arabidopsis plants used for infection assays were grown on semi-sterile soil heated in an oven at 80°C for 8 h. The plants were grown under short day conditions (8 h light and 16 h darkness) at a temperature of 22°C and a relative humidity of 65% in a growth cabinet (Percival Scientific, Perry, USA). Arabidopsis plants for seed propagation were grown under long day conditions (16 h light and 8 h darkness), light intensity of 120-150 μmol m^-2^ s^-1^, at 22°C and 60% relative humidity in a climate chamber (York Industriekälte, Mannheim, Germany). The Arabidopsis Columbia-0 (Col-0) and Wassilewskija (Ws-4) ecotypes sourced from Nottingham Arabidopsis Stock Centre were used for this work. The Arabidopsis mutants *bxl1-1* (Ws ecotype; CS16299, Feldmann, 1991), *mpk3*-DG *(Li et al., 2002)* and *dde2-2* (von Malek et al., 2002) and *B. cinerea* strain B05-10 (Staats and van Kan, 2012) were used for this work. The *Botrytis* spores were cultured on Potato dextrose broth (Sigma) plus agar, grown at RT for 10 d before harvesting by washing the spores off the plates using ¼ potato dextrose broth and sieving through Miracloth to collect the conidiospores. Conidiospores were counted using a hemacytometer (Sigma) and stocks in 25% glycerol were made and stored at −80°C. T-DNA mutant lines of *bxl4-1* (SALK_071629) and *bxl4-2* (SAIL_331_B06) were sourced from Nottingham Arabidopsis Stock Centre and homozygous mutants where confirmed through genotyping PCR on genomic DNA using REDTaq® ReadyMix™ (Sigma) following their protocol. Primers used are listed in table supplemental table 1.

### Gene expression analysis (qRT-PCR)

RNA was extracted from leaves using Spectrum™ Plant Total RNA Kit (Sigma). cDNA was made from DNaseI (Thermo Scientific, Waltham, MA, USA) treated RNA using Revert Aid™ H minus Reverse Transcriptase (Thermo Scientific). cDNA derived from leaf RNA was used for qRT-PCR using Takyon™ No Rox SYBR® MasterMix dTTP Blue (Eurogentec, Lϋttich, Belgium). Quantification of expression was done relative to *ACTIN8* (Ralhan et al., 2012). The primers used are listed in supplemental table 1.

### Molecular cloning and Arabidopsis transformation

The R4 Gateway Binary Vectors (R4pGWB; Nakagawa et al., 2008) were employed to make the constructs used in this work. The *TBA2* constructs with and without a Citrine tag were made by first amplifying the *TBA2* promoters using PCR and cloning into entry vector pDONRP4-P1R. The cDNA constructs were made by amplifying the cDNA with PCR (primers used are shown in supplemental table 1) and cloning into entry vector pDONR207. A tripartite LR reaction was performed to incorporate the *TBA2* promoter and cDNA into R4pGWB501 (modified vector) with and without Citrine tag (Nakagawa et al., 2008). The *BXL4* inducible overexpression lines were generated using pER8-GW-3’HAStrep, a plant binary gateway destination-/ 35S-inducible expression vector with a pER8-vector backbone (Zuo et al., 2000; Breitenbach et al., 2014). Arabidopsis plants were transformed by floral dipping as described (Clough and Bent, 1998) and the T1 seeds grown on MS medium supplemented with hygromycin for selection.

### Induction of BXL4 in the inducible overexpression lines

The induction of *BXL4* in the inducible *BXL4* overexpression lines was carried out by spraying 6 weeks old Arabidopsis plants with 50 µM β-estradiol in 0.01% (v/v) Tween20. Mock induction was performed by spraying the lines with 0.01% Tween20.

### Confocal microscopy

The transformed Arabidopsis T2 seeds were visualised under a confocal microscope for localisation of BXL proteins. Confocal images were recorded using confocal microscope Zeiss LSM 780 (Carl Zeiss Inc., Jena, Germany). Citrine was excited at 488 nm through a 488 nm major beam splitter (MBS). Detection of fluorophore was done at a wavelength of 514 – 530 nm.

### Botrytis cinerea infection assay

*B. cinerea* spores were diluted to 5x 10^4^spores per millilitre in ¼ Potato dextrose broth (Sigma-Aldrich) for drop inoculation assay or 2 x 10^5^ spores per millilitre in Vogel buffer (Vogel, 1956) for spray inoculation assay used for qRT-PCR analysis. The spores were pre-germinated for 4 h before inoculations were carried out. For drop inoculations, 6 µL of spore suspension in ¼ PDB was carefully placed on the adaxial side (away from the midrib) of a fully expanded rosette leaf of 6-7 weeks old Arabidopsis plants (at least 30 leaves were used from 10 independent plants). For spray inoculation, plants were sprayed until droplets began to run off the leaves (Mengiste et al., 2003). Inoculated plants were covered and grown under high humidity conditions for 72 h. Lesion diameters of drop-inoculated leaves were measured using a digital calliper and used to calculate lesion area. Spray-inoculated rosette leaves were harvested at day zero (d0 immediately after spraying) and at 72 h post inoculation (d3). For fungal DNA quantification, fungal DNA was extracted using plant/fungi DNA isolation kit (Norgen Biotek Corp) following the manufacturer’s protocol. The fungal β*-ACTIN* genomic DNA was quantified by qPCR (Ettenauer et al., 2014) and primers used are listed in supplemental table S1.

### Mucilage staining with ruthenium red

5 mg of Arabidopsis seeds were placed in 500 µL _dd_H_2_O in an Eppendorf tube before being gently shaken for 1 h on a rotary shaker. Water was gently removed and 500 µL 0.02% ruthenium red (Sigma) was added (Dean et al., 2007). Seeds were put back on the shaker for another 15 min before the ruthenium red was removed and seeds resuspended in 500 µL _dd_H_2_O added again. A droplet with stained seeds was placed on a microscopic slide and viewed under a light microscope.

### Monosaccharide analysis-mucilage

Mucilage was extracted by agitating 5 mg of seeds in water for two h on a rotary shaker. Seeds were then allowed to settle for a few minutes before 1 mL mucilage solution was removed and placed in a Duran culture tube. Mucilage solution was evaporated in a water bath at 40°C under nitrogen stream. Dry samples were hydrolysed for 1 h at 121°C using 2 M trifluoroacetic acid, before being evaporated again. 100 µL *allo*-inositol and 500 µL _dd_H_2_O were added to resuspend the hydrolysed mucilage. 20 µL sample was evaporated under a nitrogen stream before overnight derivatisation in 15 µL methoxyamide (MOX, 30mg/mL in anhydrous pyridine). The next day, 30 µL MSTFA (N-Methyl-N-(trimethylsilyl)trifluoroacetamide) were added and samples were analysed GC-MS 1-6 h after MSTFA addition.

### GC-MS analysis

Samples were analysed with a 7890B GC-System coupled to a 5977B MSD quadrupole set-up from Agilent Technologies. GC-separation was achieved on a HP-5 column (Agilent Technologies) using the following temperature gradient: 150°C for 2 min, 5 K/min gradient for 20 min, 15 K/min to a final temperature of 320°C, which was held for 3 min. For each run, 1 µL of the derivatised sample was injected. Identification of compounds was done by a combination of retention times compared to external standards and MS-spectra. Prepared mucilage samples were quantified relative to the internal standard *allo*-inositol. In parallel runs, monosaccharide standards of different concentration were used to determine response factors for area-to-molar amount conversion allowing absolute quantification.

### Monosaccharide analysis of the alcohol insoluble residue of leaves

The alcohol insoluble residue (AIR) was extracted from plant leaves (6-7 weeks old plants) grown in the dark for 48 h. The AIR was extracted as described (Gille et al., 2009). The leaves were flash frozen in liquid nitrogen before they were pulverised using mortar and pestle. The ground material was washed 2 times with 70% (v/v) ethanol, washed thrice with a chloroform:ethanol mixture (1:1 [v/v]), and lastly with acetone, before being air dried. Hot water pectin extraction (Yeoh et al., 2008) was used by shaking 2 mg of AIR in 1.4 mL _dd_H_2_O at 90°C for 2 h. Monosaccharide analysis was then carried out on AIR using the same GC-MS method used on mucilage as described above.

### Calculation of adherent mucilage volume

The shape of the seed was taken as a spheroid as described in Yu et al., 2014. The total length (2A) and width (2B) of the seed including the mucilage was measured and the volume calculated. The length of the seed alone without mucilage (2a) and the width without mucilage (2b) was measured and used to calculate the volume of the seed. The volume of the adherent mucilage was calculated by subtracting the volume of the seed alone from the total volume of the seed with mucilage using the formula: volume = 4/3 × 1/8 × length × width² (Supplemental figure 12).

### Dot-Blot assay

The dot blot assay was performed as described in Bethke et al., (2016). Pectin was extracted using a pectin extraction buffer (50 mM Trizma and 50 mM CDTA, pH 7.2) at 50 µL per mg AIR. Serial dilutions were done before dropping 1 µL on nitrocellulose membranes. Overnight drying of the membrane was done before the membranes were blocked by adding 5% milk powder (w/v) dissolved in 1× PBS (8 g L^−1^ NaCl, 0.2 g L^−1^ KCl, 1.44 g L^−1^ Na_2_HPO_4_, and 0.24 g L^−1^ KH_2_PO_4_, pH 7.4). The membranes were probed with LM13 (Verhertbruggen et al., 2009), LM19 (Verhertbruggen et al., 2009) and CCRC-M7 (Steffan et al., 1995) antibodies. LM antibodies were diluted 1:250 and CCRC-M7 diluted 1:500 in 5% milk powder (w/v) dissolved in 1 x PBS. Rabbit anti-rat IgG antibody (Sigma) diluted 1:30000 in 5% milk powder (w/v) in 1 x PBS was used for the LM antibodies. Goat anti-mouse IgG antibody (Sigma) diluted 1:30000 in 5% milk powder (w/v) in 1 x PBS was used for CCRC-M7 antibodies. Blots were developed by equilibrating in AP buffer (100 mM Tris, 100 mM NaCl, 5 mM MgCl_2_, pH 9.5) before incubating in 10 mL AP buffer with 33 µL BCIP and 66 µL NBT in the dark until spots were visible.

### Phytohormone measurements

Extraction of phytohormones, separation and analysis was carried out as described in Kusch et al., (2019). Mass transitions were used as described by Iven et al., (2012), with some modifications specified in supplemental table 2.

## Supplemental Data

Supplemental figure 1: Expression of *BXL4* in Arabidopsis rosette leaves.

Supplemental figure 2: Phylogenetic tree of betaxylosidases (BXLs) from *Arabidopsis thaliana*.

Supplemental figure 3: *bxl4* mutants show normal extrusion of mucilage.

Supplemental figure 4: BXL1-CITRINE and BXL4-CITRINE localise to the apoplast in Arabidopsis *bxl1* seed coat epidermal cells

Supplemental figure 5: BXL4 without a CITRINE tag complements the mucilage phenotype of *bxl1*.

Supplemental figure 6: BXL6 fails to complement the mucilage phenotype of *bxl1*.

Supplementary figure 7: Mucilage monosaccharide composition

Supplemental figure 8: Monosaccharide composition of pectin extracted from leaf AIR

Supplemental figure 9: Bot blot analysis on pectin extracted from leaf alcohol insoluble residue (AIR).

Supplemental figure 10: Morphological phenotypes of Col-0, *bxl4-1* and *bxl4-2*

Supplemental figure 11: BXL4 acts upstream of jasmonoyl-isoleucine signalling upon *B. cinerea* infection.

Supplemental figure 12: JA-Ile accumulation after mechanical wounding of Arabidopsis leaves.

Supplemental figure 13: Calculation of adherent mucilage volume

Supplemental table 1: List of primers used

Supplemental Table 2: Mass transitions and corresponding conditions for determination of the phytohormones.

Supplemental dataset1: Phytohormone analysis

## AUTHOR CONTRIBUTIONS

A.G., C.V., G.H., M.W., and T.I. designed the research. A.G., R.M., D.H., P.S., K.B., MW., D.L. and K.Z. performed the experiments. M.W., C.V., I.F., G.H., and T.I. provided the tools and materials and analysed the data. A.G. and T.I wrote the manuscript with contributions from the other authors.

## ACKNOWLEDGEMENTS

This work was supported by German Research Foundation (DFG, IRTG 2172 PRoTECT to T.I., M.W., and I.F.), the Studienstiftung des deutschen Volkes (stipend to P.S.), a University of British Columbia four-year fellowship to R.M. and a Natural Sciences and Engineering Research Council of Canada Discovery grant to G.W.H. We would like to thank Prof. Dr. Volker Lipka for useful discussions and suggestions to improve the manuscript, Franziska Kretzschmar for proofreading the manuscript. We are also grateful to the following individuals who helped with work in the lab: Maurice Hädrich, Maximillian Sievers, Finni Häußler and Milena Lewandowska. We would also like to thank Prof. Großhans and Prof. Johnsen together with Dr. Florian Wegwitz and Johannes Sattmann for granting us access to the microscopes and their assistance.

