## Supplemental figures 1-11 for "The cell-wall-localised BETA-XYLOSIDASE 4 contributes to immunity of Arabidopsis against *Botrytis cinerea*"

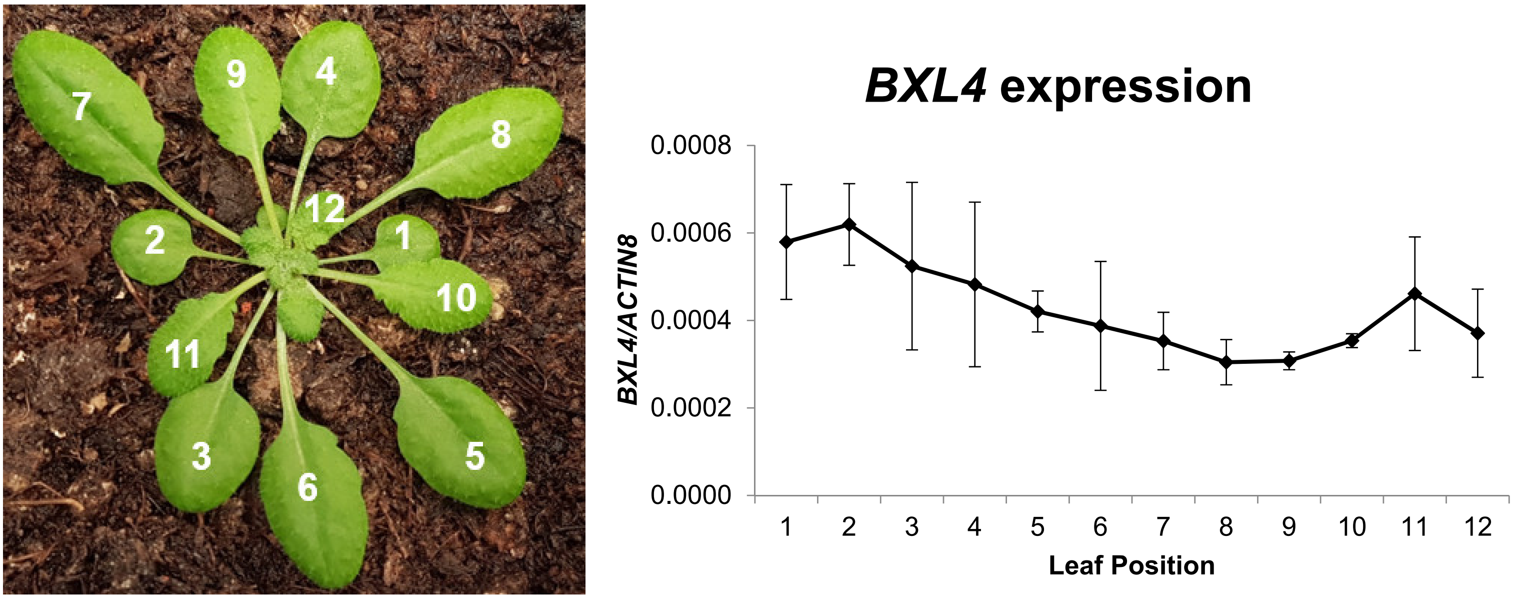


**Supplemental figure 1: Expression of *BXL4* in Arabidopsis rosette leaves.**
RNA was extracted from the different Arabidopsis (Col-0) leaves starting from the oldest rosette leaf (1) to the youngest (12). Expression of *BXL4* was determined by qRT-PCR and normalised to the expression of *ACTIN8*. Error bars represent standard error of three biological replicates.

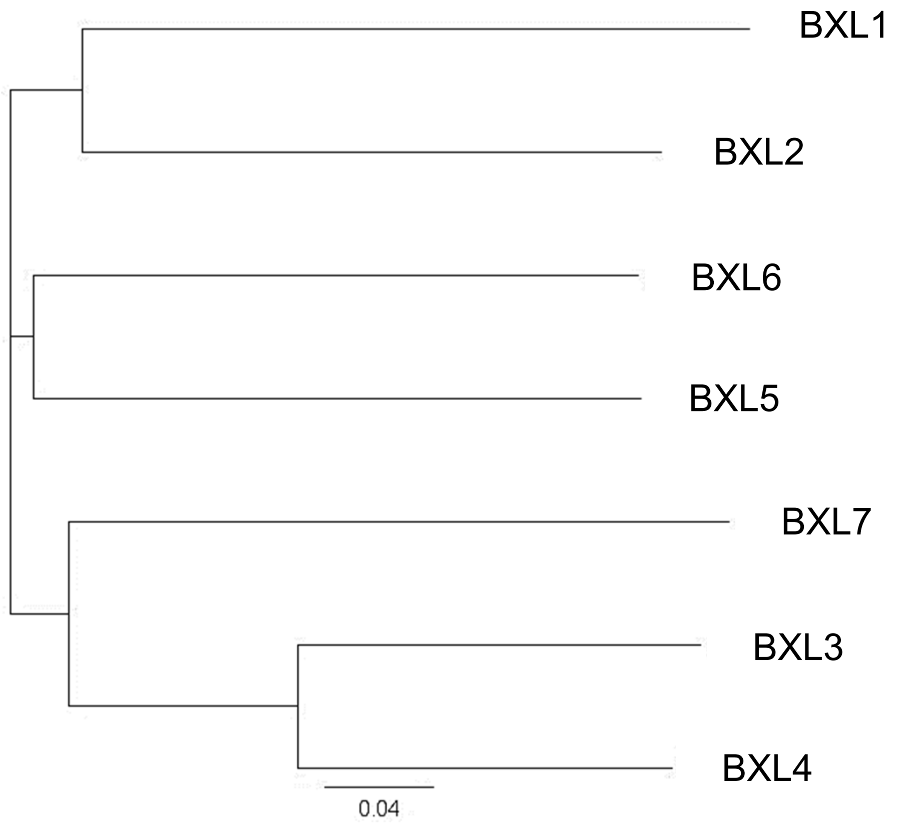


**Supplemental figure 2: Phylogenetic tree of betaxylosidases (BXLs) from *Arabidopsis thaliana*.** Image generated with Geneious version 8.1 using Jukes-Cantor genetic distance model and the Neighbour-joining tree method.

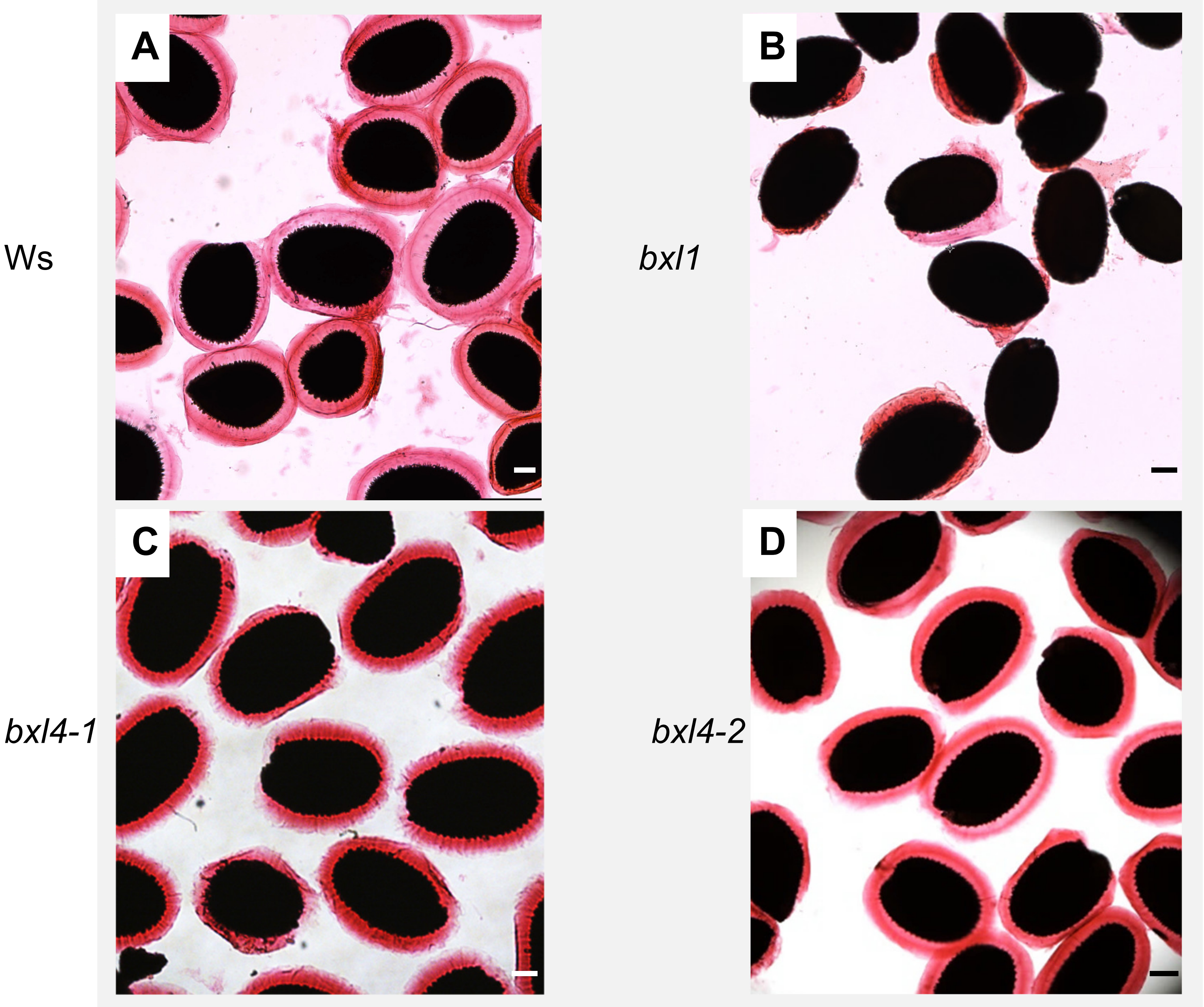


**Supplemental figure 3: *bxl4* mutants show normal extrusion of mucilage.**
The extrusion of the ruthenium red-stained mucilage from Ws, *bxl1*, *bxl4-1* and *bxl4-2* (**A**, **B**, **C** and **D** respectively**)**. Scale bars, 100 µm.

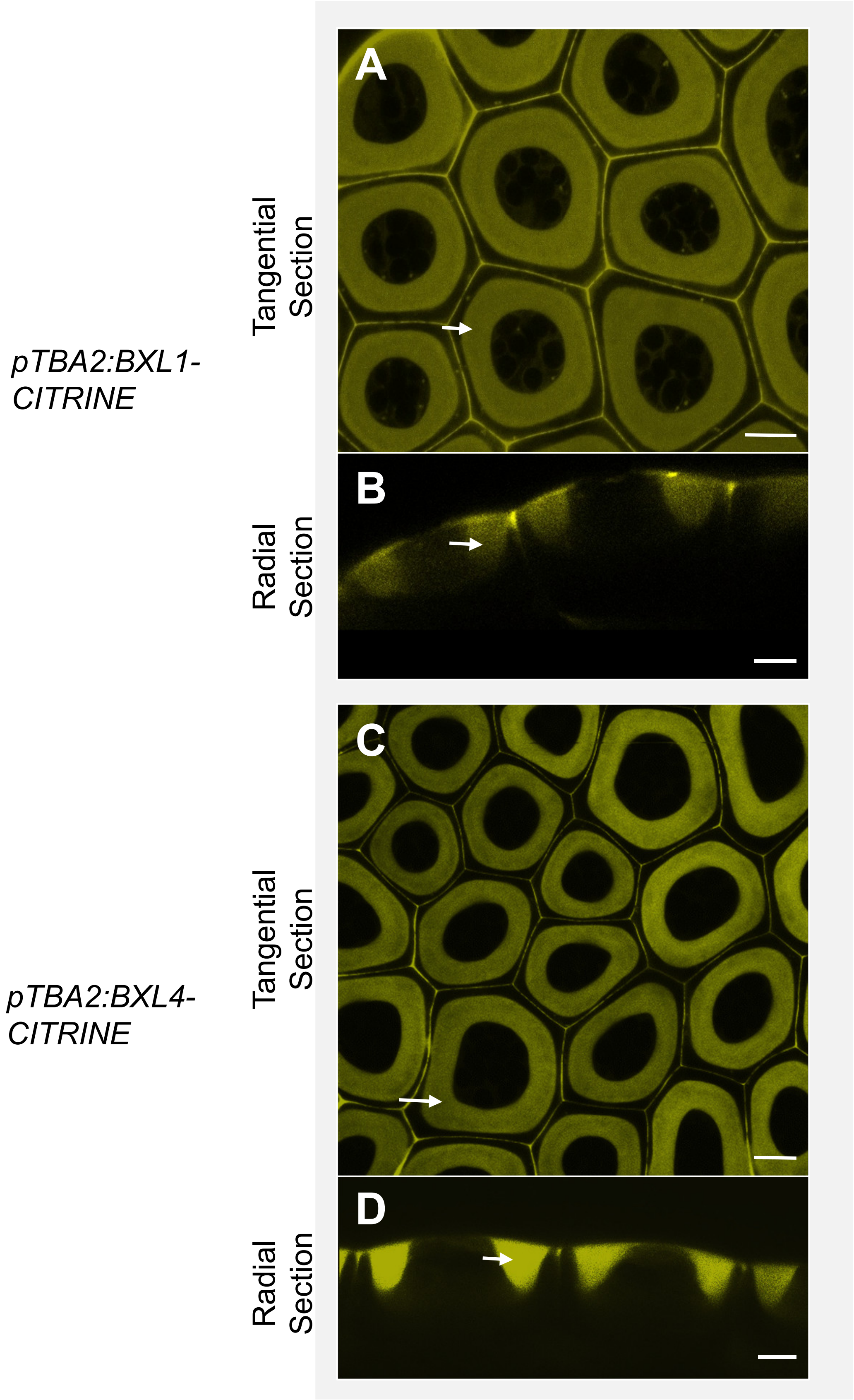


**Supplemental figure 4: BXL1-CITRINE and BXL4-CITRINE localise to the apoplast in Arabidopsis *bxl1* seed coat epidermal cells**

**(A)** BXL1-CITRINE stably expressed under control of the TBA2 promoter localises to the apoplast (deposited predominantly in the mucilage pocket) of Arabidopsis *bxl1* seed coat epidermal cells at 7 days post anthesis. Single plane images of the seed coats were obtained by confocal microscopy. The apopolast of seed coat epidermal cells appears like a doughnut ring (arrow) that surrounds the cytoplasm when imaged tangentially. **(B)** Visualisation of the seed coat epidermal cells radially shows the apoplast as two pockets in the apical corners of the cells (arrow). BXL4-CITRINE expressed by the TBA2 promoter also localises to the apoplast of Arabidopsis seed coat epidermal cells **(C, D)**. Scale bars, 10 µm.

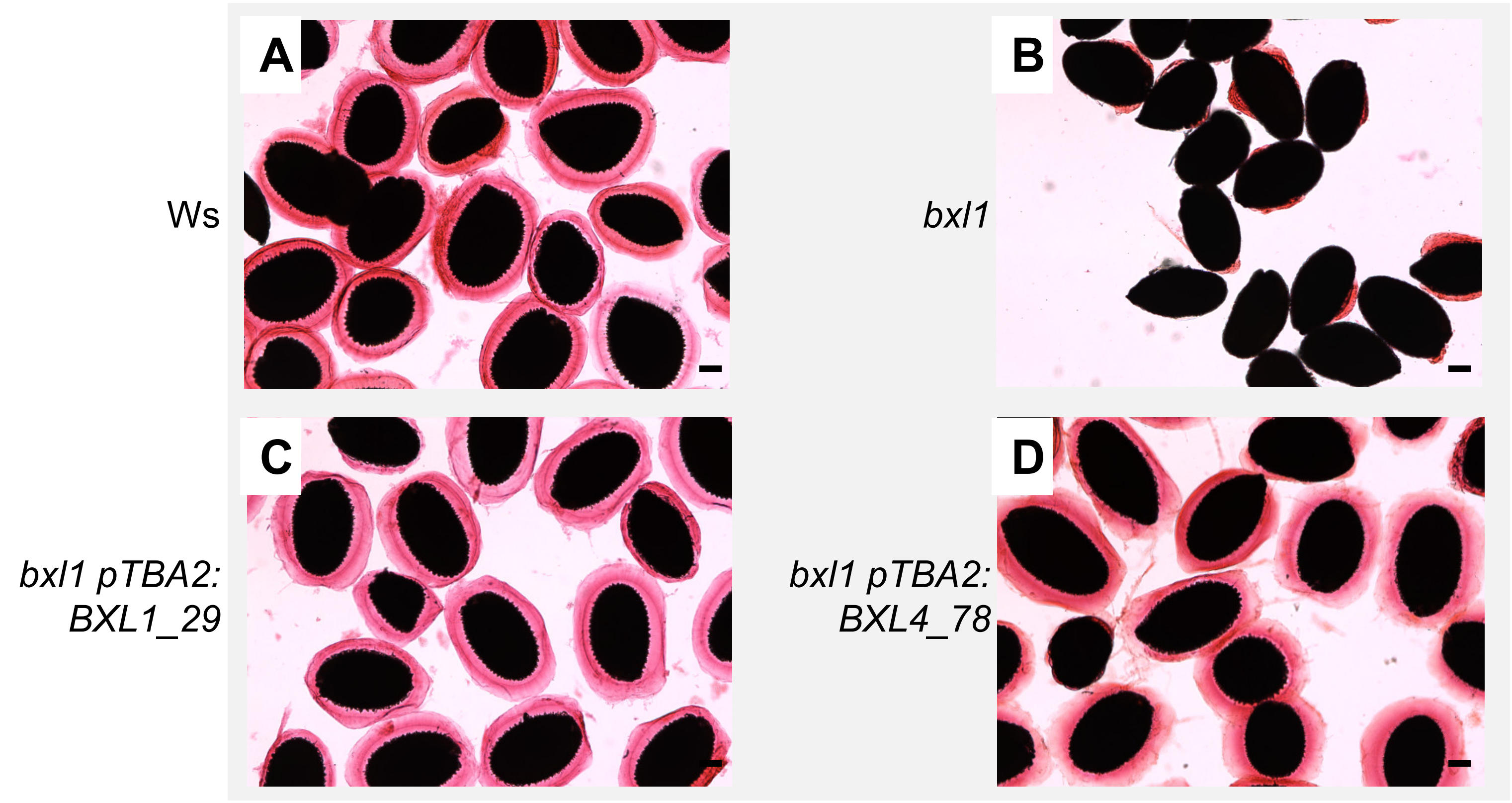


**Supplemental figure 5: BXL4 without a CITRINE tag complements the mucilage phenotype of *bxl1*.** Seeds of Ws **(A)**, *bxl1* **(B)**, *bxl1* transformed with *pTBA2:BXL4* **(C)** *bxl1* transformed with *pTBA2:BXL4* **(D)** hydrated and stained with ruthenium red.

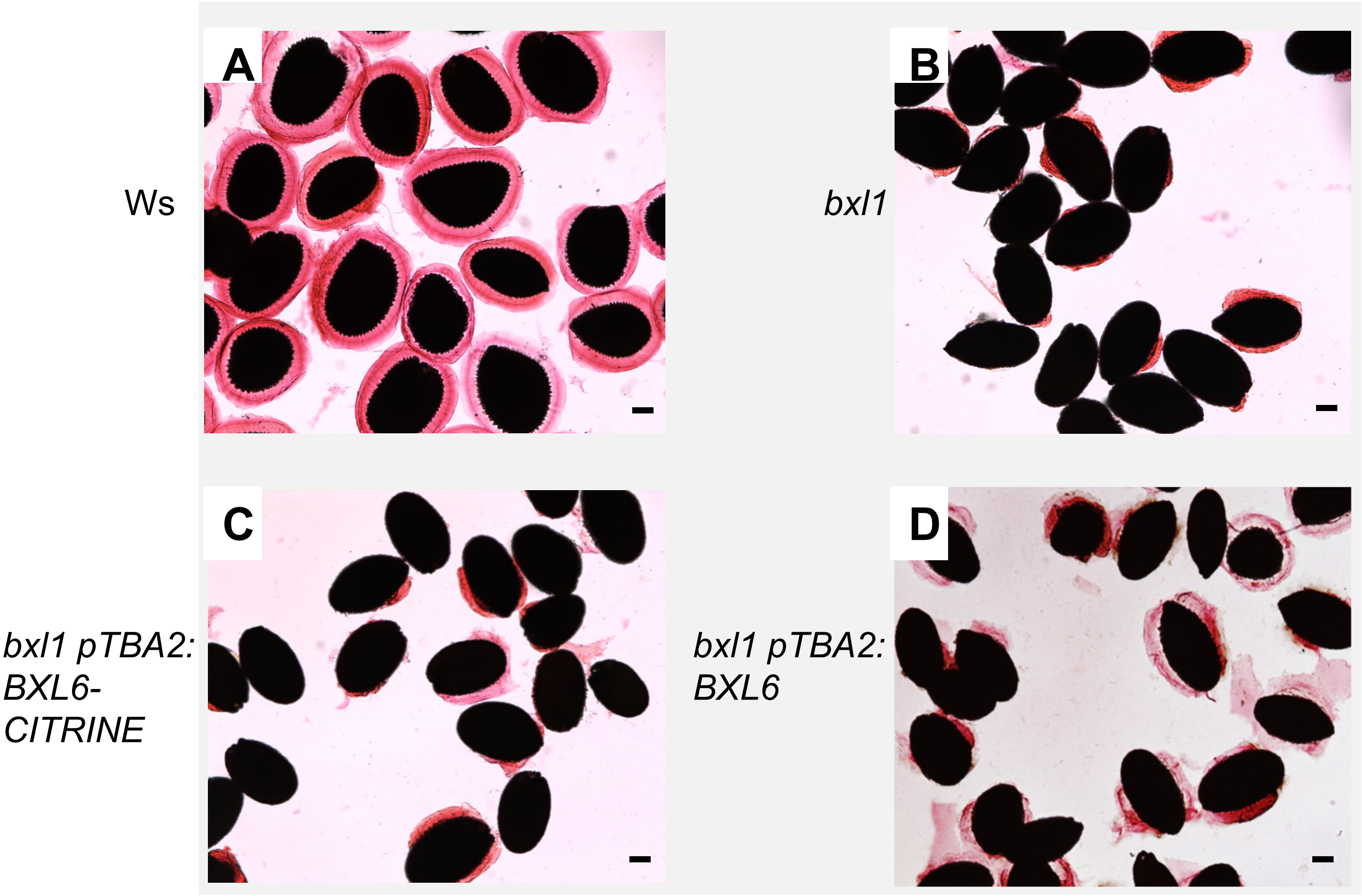


**Supplemental figure 6: BXL6 fails to complement the mucilage phenotype of *bxl1*.**
Transgenic *bxl1* plants expressing *pTBA2:BXL6* fail to extrude their mucilage normally. Seeds of Ws **(A)**, *bxl1* **(B)** and two independent *bxl1* transgenic lines expressing *pTBA2:BXL6-CITRINE* **(C)** *pTBA2:BXL6* **(D)** hydrated and stained with ruthenium red. Scale bars, 100 µm

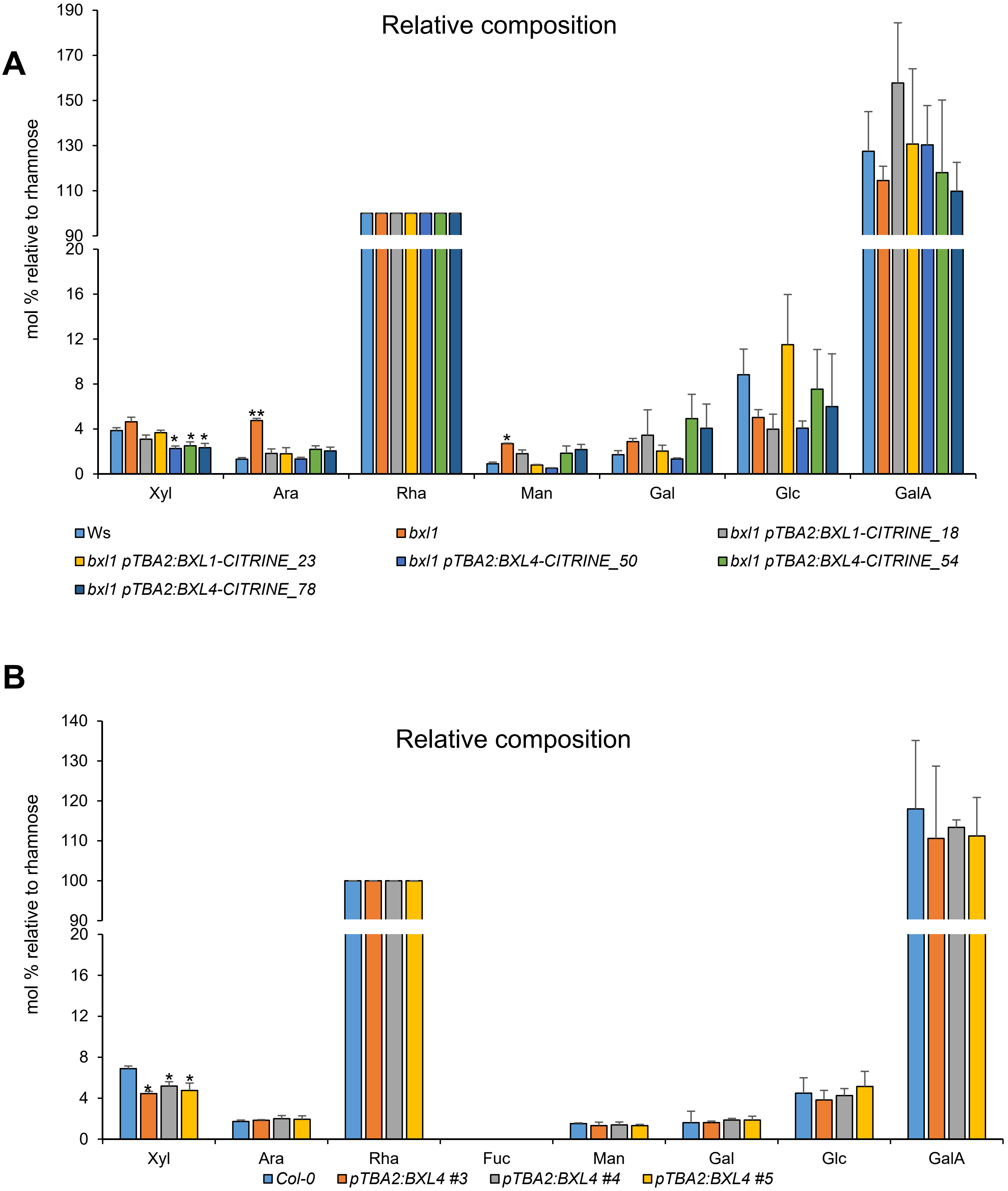


**Supplementary figure 7: Mucilage monosaccharide composition**
**(A)** Relative monosaccharide composition of mucilage extracted from Ws**,** *bxl1, bxl1* p*TBA2*:*BXL1-CITRINE* (line 18 and 23) and *bxl1* p*TBA2*:*BXL4-CITRINE* (line 50, 54 and 78).
**(B)** Monosaccharide composition of mucilage extracted from wild type Col-0 and three *BXL4* overexpression lines (p*TBA2*:BXL4 line 3, 4 and 5). Monosaccharide composition was determined by GC-MS and normalised to rhamnose. n=3 biological replicates. n=3 biological replicates. Error bars show SD, statistical difference to WT (Student's *t*-test ),* indicates p<0.05, ** p<0.01

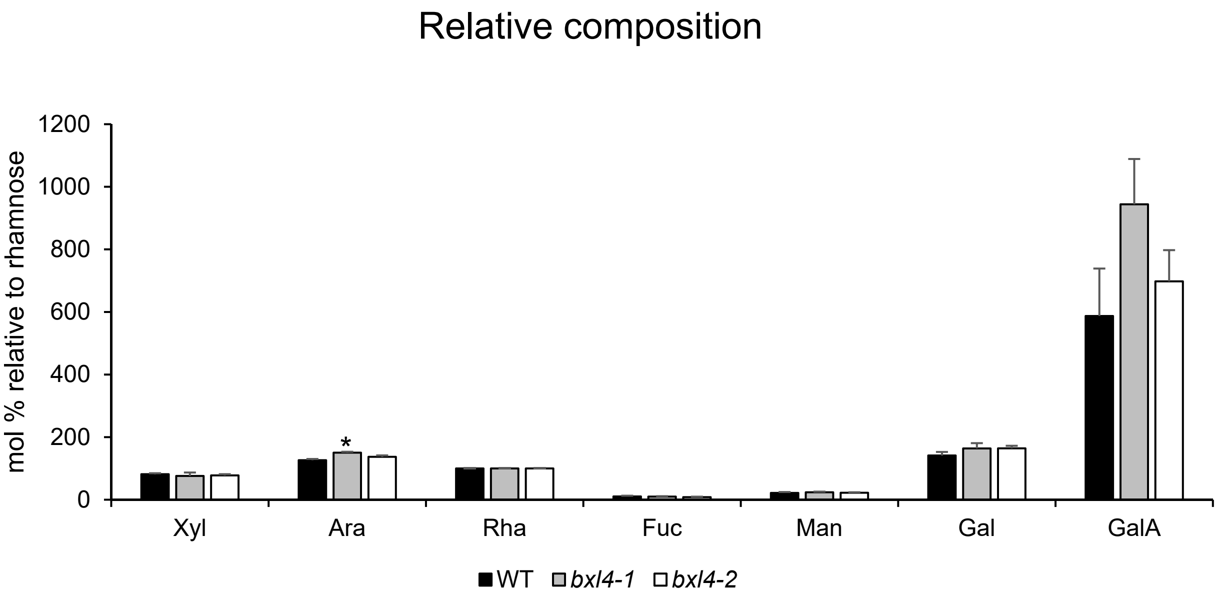


**Supplemental figure 8: Monosaccharide composition of pectin extracted from leaf AIR.**

Monosaccharide composition of water extracted pectin from Arabidopsis leaves of wildtype, *bxl4-1* and *bxl4-2* mutant lines. The monosaccharides were normalised to rhamnose. Extracted pectin was analysed by GC-MS. Error bars show SD (n=4 biological replicates), statistical difference to WT (Student's *t*-test ), *p<0.05. Experiments were done three times with similar results

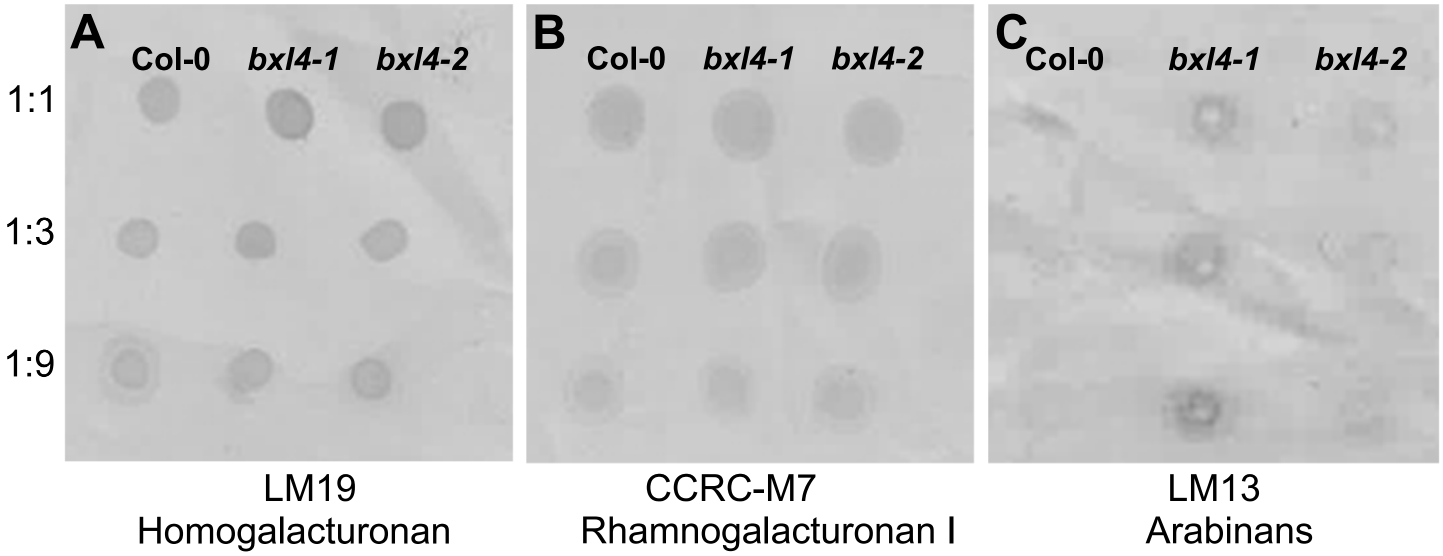


**Supplemental figure 9: Dot blot analysis on pectin extracted from leaf alcohol insoluble residue (AIR).**
Pectin was extracted from the AIR of the three indicated genotypes using pectin extraction buffer. The pectin was serially diluted before 1 µL was spotted onto a nitrocellulose membrane. The dot blot was carried out using LM19 **(A)**, CCRC-M7 **(B)** and LM13 **(C)**.

3 weeks

**A**

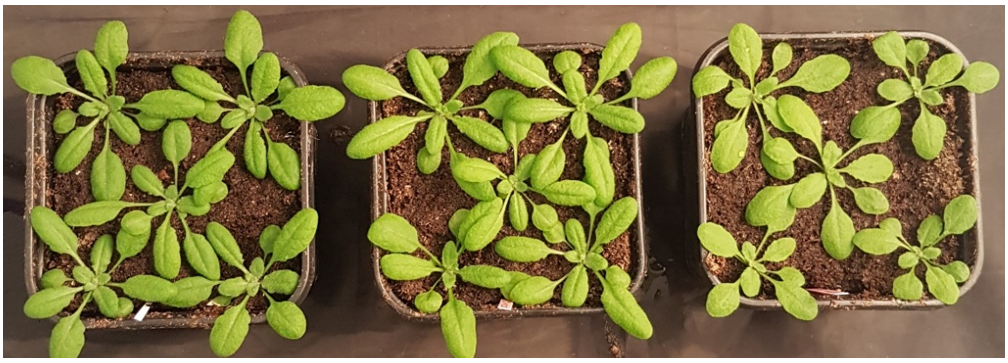

8 weeks

**B**

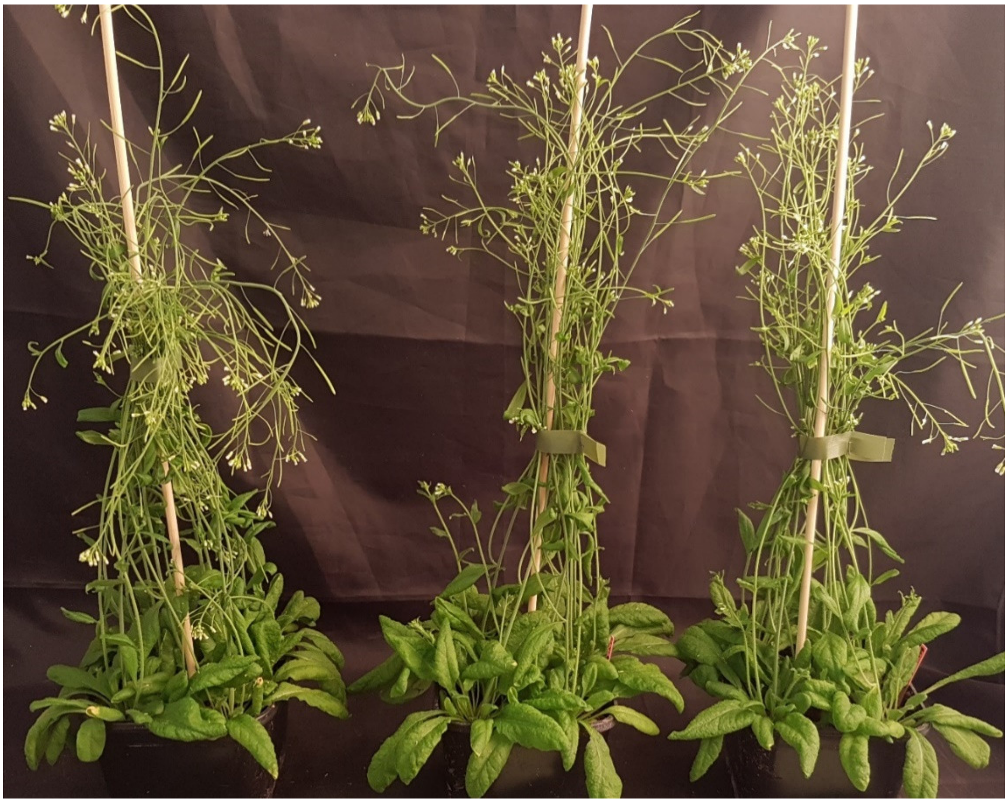

**Col-0** ***bxl4-1*** ***bxl4-2***

**Supplemental figure 10: Morphological phenotypes of Col-0, *bxl4-1* and *bxl4-2*.** The morphological phenotypes of 3 weeks **(A)** and 8 weeks old **(B)** Col-0, *bxl4-1* and *bxl4-2* grown under long day conditions (16 h light/8 h dark).

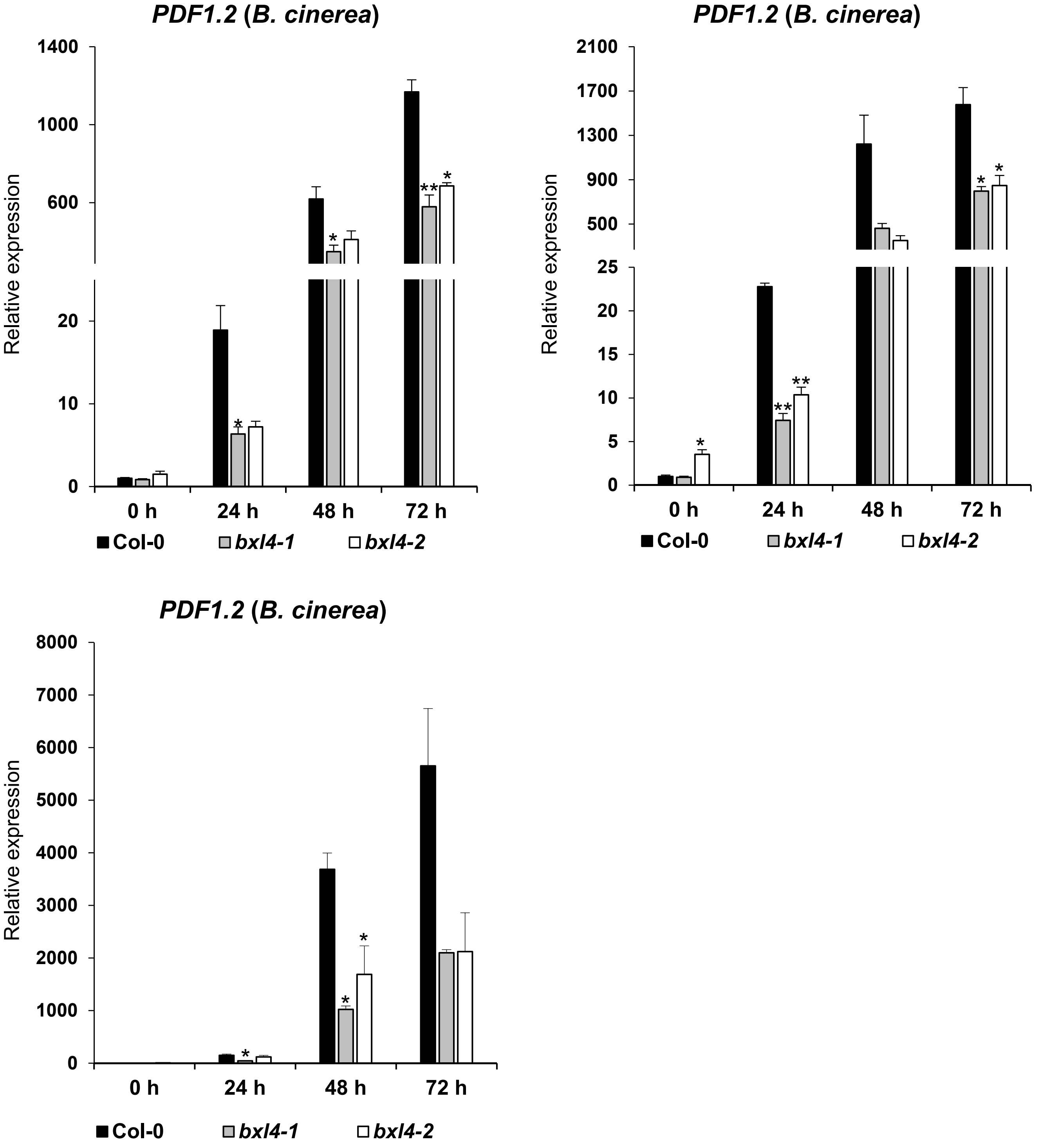


**Supplemental figure 11: BXL4 acts upstream of JA-Ile signalling upon *B. cinerea* infection.** Relative expression of *PDF1.2* in RNA extracted from 6 week old Col-0, *bxl4-1* and *bxl4-2* Arabidopsis plants at 0, 24, 48 and 72 h after infection with *B. cinerea.* Data shown is from 3 independent experiments **(A-C).** Expression values were normalised to the reference gene *ACTIN8* and are shown relative to the wild type 0 h levels. Error bars show SE (n=3 biological replicates), statistical difference to WT (Student's *t*-test ), *p<0.05, **p<0.01.

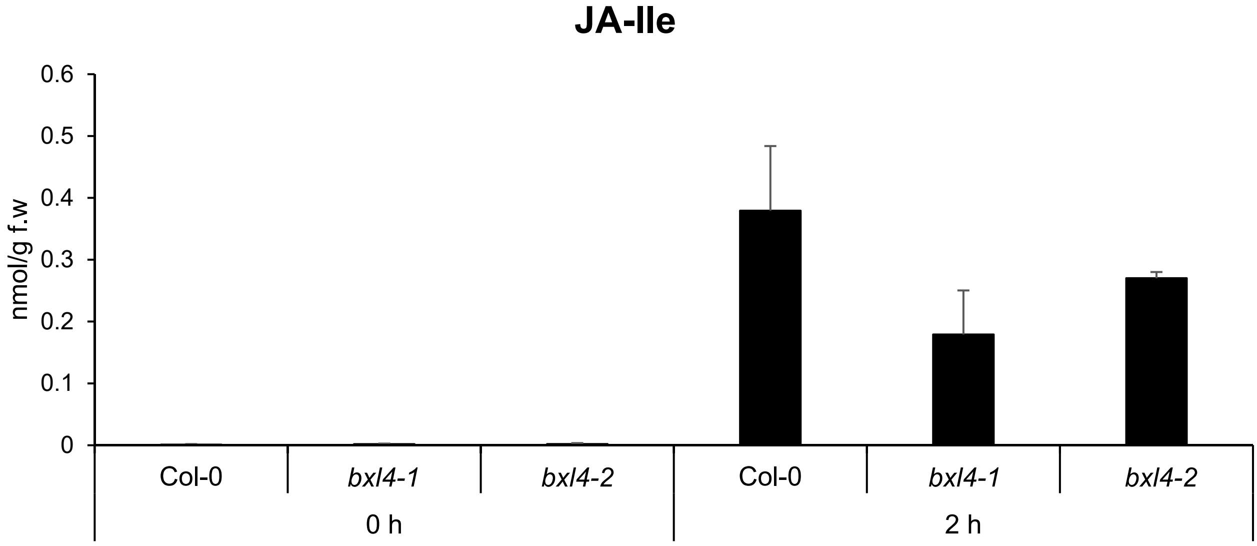


**Supplemental figure 12: JA-Ile accumulation after mechanical wounding of Arabidopsis leaves.**
The leaves of Col-0 and *bxl4* mutant lines were mechanically wounded and sampled at 0, 2, 24 h after wounding. Extracted levels of JA-Ile were analysed using nanoelectrospray coupled to a tandem mass spectrometer. Error bars represent standard deviation of 3 biological replicates

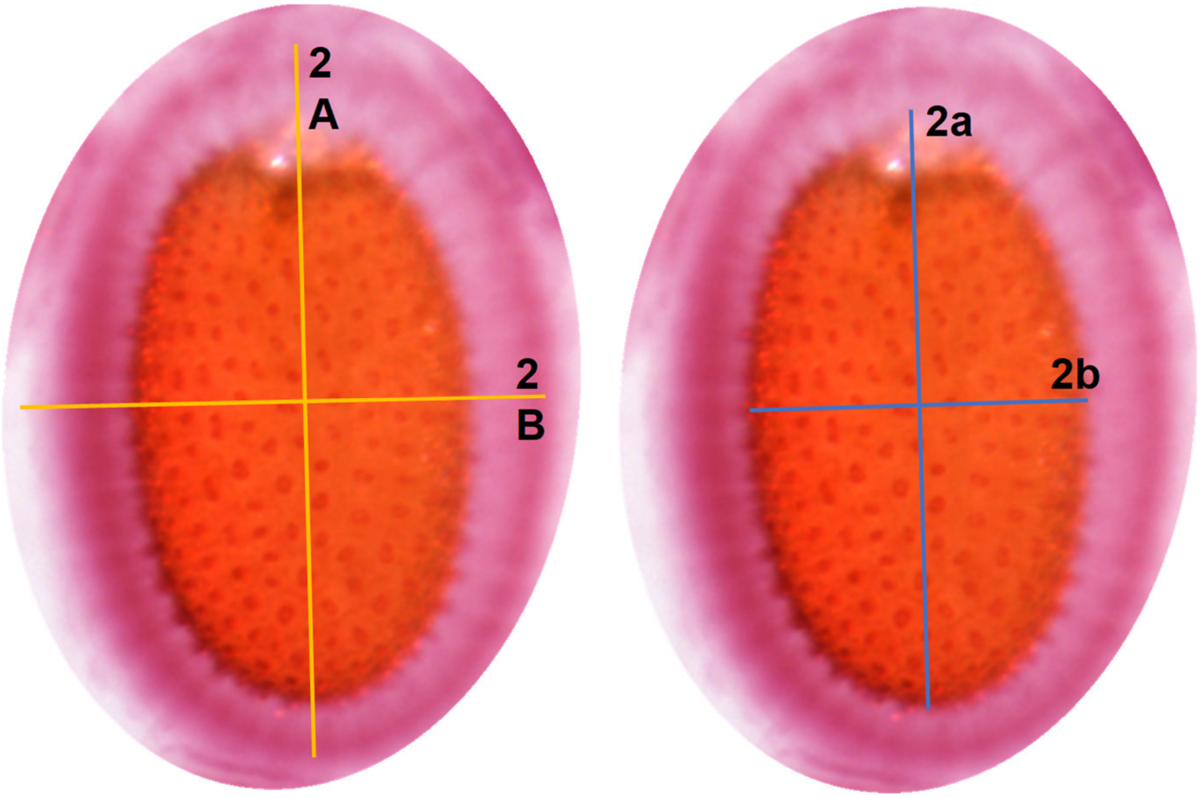


**Supplemental figure 13: Calculation of adherent mucilage volume**

The shape of the seed was taken as a spheroid as described in Yu et al., (2014). The total length 2A and width (2B) of the seed including the mucilage was measured and the volume calculated. The length of the seed alone without mucilage (2a) and the width without mucilage (2b) was measured and used to calculate the volume of the seed. The volume of the adherent mucilage was calculated by subtracting the volume of the seed alone from the total volume of the seed with mucilage using the formula: volume = 4/3 × 1/8 × length × width × depth. The width of the seed was used as the depth.
