## Supplemental figures 1-2 for "The cell-wall-localised BETA-XYLOSIDASE 4 contributes to immunity of Arabidopsis against *Botrytis cinerea*"

**Supplemental table 1: List of primers used**

| **Cloning primers** | **Forward 5’-3’** | **Reverse 5’-3’** |
|---|---|---|
| *BXL1* | GGGGACAAGTTTGTACAAAAAAGCAGGCTATGTCTTGTTATAATAAAGCACTATTG | GGGGACCACTTTGTACAAGAAAGCTGGGTAAAGTTGCGGTTGGACCAA |
| *BXL4* | GGGGACAAGTTTGTACAAAAAAGCAGGCTTGGGCTCTTCTTCTCCATTA | GGGGACCACTTTGTACAAGAAAGCTGGGTAGATTCTAATGCTTAAGGAATGTTTTA |
| *BXL6* | GGGGACAAGTTTGTACAAAAAAGCAGGCTATGAATCTTCAGTTGACTCTAATC | GGGGACCACTTTGTACAAGAAAGCTGGGTAGAATTCAACAGAGAGAGAATGT |
| *BXL4* (OE) | CACCATGGGCTCTTCTTCTCC | GATTCTAATGCTTAAGGAATGTTTTAAATCTCCG |
| **Genotyping primers** | **LB** | **RB** |
| *bxl1* | ATTTTGCCGATTTCGGAAC (LBb1.3) | AACCGTCGCGTCGGCTTCAC |
| *bxl4-1* | LBb1.3 | ATCTCCGACATGAAGAAGATGC |
| *bxl4-2* | TAGCATCTGAATTTCATAACCAATCTCGATAC | ATCTCCGACATGAAGAAGATGC |
| *bxl6* | LBb1.3 | TACCACAGCATTGAAGTCGTATC |
| **qRT-PCR primers** | **Forward 5’-3’** | **Reverse 5’-3’** |
| *BXL4* | ATACACAACACCACTACAAGGAC | ATCTCCGACATGAAGAAGATGC |
| *BXL4a* | TCAACGCCGTGGTGACTCAA | CGCATGTCGGTTTGCCGTTA |
| *BXL4b* | CCCACACCTGTTTTCAGTGCC | TACATTGCCCTCGCTTCCGT |
| *PDF1.2* | TTGCTGCTTTCGACGCA | TGTCCCACTTGGCTTCTCG |
| *JAZ10* | ATCCCGATTTCTCCGGTCCA | ACTTTCTCCTTGCGATGGGAAGA (Benthke et al., 2016) |
| *PAD3* | TGCTCTCAAGTTCACCACT | CGAATCTCGTCTTGCACTT (Benthke et al., 2016) |
| *ACTIN8* | GGTTTTCCCCAGTGTTGTTG | CTCCATGTCATCCCAGTTGC (Ralhan et al., 2012) |
| *Botrytis ACTIN* | TGGAGATGAAGCGCAATCCA | AAGCGTAAAGGGAGAGGACG |
| *Botrytis TUBULIN* | CCGTCATGTCCGGTGTTAC | CGACCGTTACGGAAATCGG |

**Supplemental Table 2: Mass transitions and corresponding conditions for determination of the phytohormones.**

| MRM Transitions | | Analyte | DP [declustering potential] | EP [entrance potential] | CE [collision energy] |
|---|---|---|---|---|---|
| Q1 | Q3 | | | | |
| 209 | 59 | JA | -30 | -4.5 | -24 |
| 225 | 59 | 11,12-0H-JA | -35 | -9 | -28 |
| 263 | 165 | dinor-oPDA | -40 | -5 | -20 |
| 296 | 170.2 | D5-oPDA | -65 | -4 | -28 |
| 305 | 97 | 12-HS04-JA | -30 | -4 | -32 |
| 308 | 116 | JA-Val | -45 | -5 | -28 |
| 322 | 130 | JA-lle/Leu | -45 | -5 | -28 |
| 325 | 133 | D4-JA-Leu | -80 | -4 | -30 |
| 324 | 116 | 120H-JA-Val | -45 | -10 | -30 |
| 338 | 130 | 120H-JA-lle | -45 | -10 | -30 |
| 352 | 130 | 12COOH-JA-lle | -45 | -10 | -30 |
| 387 | 59 | 12-0-Gluc-JA | -85 | -9 | -59 |
